# Cardiomyocyte-specific expression of HIF-1α mediates the cardioprotective effects of Growth Hormone Releasing Hormone (GHRH)

**DOI:** 10.1101/2025.07.03.662389

**Authors:** Rosemeire M. Kanashiro-Takeuchi, Lauro M. Takeuchi, Wayne Balkan, Amarylis C.B.A. Wanschel, Konstantinos E. Hatzistergos, Shathiyah Kulandavelu, Russell G. Saltzman, Lina A. Shehadeh, Wei Sha, Andrew V. Schally, Stefan Kurtenbach, Joshua M. Hare

**Affiliations:** Interdisciplinary Stem Cell Institute, University of Miami Miller School of Medicine, Miami, FL 33136, USA; Department of Molecular and Cellular Pharmacology, University of Miami Miller School of Medicine, Miami, FL 33136, USA; Department of Medicine, Division of Cardiovascular Medicine, University of Miami Miller School of Medicine, Miami, FL 33136, USA; Department of Cell Biology, University of Miami Miller School of Medicine, Miami, FL 33136, USA; Department of Genetics, Development and Molecular Biology, Aristotle University of Thessalonik, 54124 Thessaloniki, Greece; Department of Pediatrics, Division of Pediatric Nephrology, University of Miami Miller School of Medicine, Miami, FL 33136, USA; Veteran Affairs Medical Center, FL 33125, USA; Department of Pathology, University of Miami Miller School of Medicine, Miami, FL 33136, USA

**Keywords:** Growth Hormone Releasing Hormone, HFpEF, HIF-1α, Cardiac-Specific Cre

## Abstract

Heart failure (HF) with preserved ejection fraction (HFpEF) carries a high mortality and remains a major therapeutic challenge. Effective, targeted therapies capable of reversing HFpEF pathophysiology are urgently needed. We previously demonstrated that activation of the cardiac growth hormone-releasing hormone (GHRH) pathway using high potency synthetic agonists of GHRH (GHRH-agonists: MR-356 and MR-409) improves the HFpEF phenotype in both large and small animal models, including in a murine model of cardiometabolic HFpEF (High fat diet + the nitric oxide synthase inhibitor L-NAME [HFD+L-NAME]). Here we sought to define the downstream signaling pathways responsible for this effect. A transcriptomic screen in human iPSC-derived cardiomyocytes (hiPSC-CMs) identified the hypoxia-inducible factor (HIF)-1α pathway as being activated by GHRH receptor (GHRHR) signaling, revealing an oxygen-independent mechanism of HIF-1α activation. Based on this finding, we investigated the interaction between the cardioprotective effects of the GHRH-agonist MR-356 and HIF-1α pathway activation in the murine HFD+L-NAME model of cardiometabolic HFpEF. We generated a cardiomyocyte-specific HIF-1α knockout (HIF-1α^CM^KO) mouse line and demonstrated that our previously reported beneficial effects of GHRH-agonist administration were completely abolished in HIF-1α^CM^KO mice. Together, these findings establish the GHRHR-HIF-1α axis as a central pathway integrating metabolic and contractile remodeling, suggesting that therapeutic targeting of this axis represents a novel disease-modifying approach to treating cardiometabolic HFpEF.

## Introduction

Heart failure with preserved ejection fraction (HFpEF) has a multifactorial pathophysiology and presents substantial clinical heterogeneity^1, 2^. Cardiometabolic HFpEF is expected to become the most prevalent form of HF^3–5^. Treatments are limited, and they only ameliorate but do not reverse disease pathophysiology. Therefore, effective management strategies and novel interventions specifically targeting the pathophysiological mechanisms underlying HFpEF are essential for advancing the development of new therapies.

Growth hormone-releasing hormone (GHRH) provides benefits in both preclinical and clinical studies for the cardiovascular system^6^, aging, and obesity^7^. Various studies have shown therapeutic effects of GHRH agonists in animal models of acute^8–10^ and chronic^11^ ischemic heart diseases and, more recently, in three distinct animal models of HFpEF (murine Angiotensin II-induced^12^, cardiometabolic^13^ HFpEF, and a swine model of cardiorenal HFpEF^14^). However, the mechanisms and signaling pathways underlying these cardioprotective effects remain unclear. Here, we show that activating the GHRH receptor (GHRHR) upregulates hypoxia-induced factor (HIF)-1α in iPSC-derived cardiomyocytes. HIF-1α is a transcription factor that functions as a master regulator of cardiac oxygen homeostasis, including the regulation of oxygen utilization through glucose metabolism and redox homeostasis^15–17^, and is an oxygen-sensitive regulator of cardiac protection (reviewed in^18^). In this study, we aimed to elucidate the molecular mechanisms of cardioprotection through GHRHR activation and HIF-1α signaling, so as to provide a fuller understanding of the cellular machinery that drives HFpEF and yield more effective therapeutic strategies. In addition, we confirmed the essential role of HIF-1α in mice with a cardiomyocyte (CM)-selective deletion of HIF-1α (HIF-1α^CM^KO) and interrogated the effects of GHRH agonists on the reversal of the adverse impacts of HFpEF on the structural, functional and metabolic remodeling of the heart.

## Methods

All experimental procedures were reviewed and approved by the University of Miami Institutional Animal Care and Use Committee (Protocol # 21-044) and comply with all Federal and State guidelines concerning the use of animals in research and teaching as defined by *The Guide for the Care and Use of Laboratory Animals* (revised 2011)^19^. Mice were housed in an AAALAC-accredited animal facility at the University of Miami Miller School of Medicine Division of Veterinary Resources (DVR). Their care was supervised by licensed veterinarians. The University of Miami has an Animal Welfare Assurance on file with the Office of Laboratory Animal Welfare (OLAW), National Institutes of Health. The assurance number is #A-3224-01, valid until November 30, 2027. UM has received full accreditation with the Association for Assessment and Accreditation of Laboratory Animal Care (AAALAC International), site 001069, latest effective date of June 6, 2022.

### Human iPSC-derived cardiomyocytes (hiPSC-CMs)

The culture and differentiation of hiPSCs (SC101A, System Biosciences) were conducted as previously described^20^ with minor modifications. Briefly, hiPSCs were maintained in E8 medium on Matrigel-coated plates. They were dissociated into single cells using TrypLE (GIBCO) and seeded at a density of 1 × 10^5 cells per well in a 12-well plate. The cells were grown for 96 hours until reaching 90% confluence in a humidified incubator at 37°C with 5% CO2. On day 0, the medium was replaced with RPMI 1640 supplemented with B27 (without insulin) and 6 μM CHIR99021 (2520691, Biogems). On day 1, the medium was changed to RPMI 1640 with B27 (without insulin), and 5 μM IWP2 (6866167, Biogems) was added after 24 hours, continuing for 72 hours. On day 4, the medium was switched to RPMI 1640 supplemented with 3 μg/mL heparin, with medium changes every 2 days until day 8. From day 8 onwards, fresh RPMI 1640 containing 3 μg/mL heparin and 20 μg/mL insulin was changed every 2 days. For GHRH induction experiments, wild-type (WT) and HIF1α-KO cells were treated with 0, 0.15, or 0.3 μM recombinant human GHRH (ab52521, Abcam) for 45 minutes (n=3/genotype/dose).

### Quantitative PCR

Total RNA was extracted using the RNeasy Mini Kit (74106, Qiagen), and complementary DNA (cDNA) was synthesized with the High-Capacity cDNA Reverse Transcription Kit (4368814, Applied Biosystems). Quantitative PCR was performed using an iQ5 real-time PCR detection system (Bio-Rad) with TaqMan Universal Master Mix (Applied Biosystems). The following probes were used: GHRH-R (Hs0181591), GHRH (Hs00184139), and HIF-1α (Hs00153153).

### Western blotting

Whole cell protein was extracted using the Active Motif Protein Extraction Kit and quantified by the Bradford assay (Bio-Rad). Electrophoresis was performed in precast NuPage 4-12% Bis-Tris protein gels (NP0323, Thermo Scientific; MP42G15, Merck) before transferring to 0.22μm polyvinylidene difluoride (PVDF) membranes (LC2002, Thermo Fisher Scientific) using the TransBlot Turbo (Bio-Rad) or Mini Blot Module (B1000, Thermo Fisher Scientific). Western blots were conducted following blocking with 3% Blotto (2325, Santa Cruz Biotechnology) for 40 minutes. Primary antibodies included: HIF-1α (1:2000, D1S7W, Cell Signaling), GHRH-R (1:500, ab76263, Abcam for mouse tissue), GHRH-R (1:1000, LS-C383690, LSBio for human cells), β-actin (1:6000, 8H10D10, Cell Signaling), GAPDH (1:2000, D16H11, Cell Signaling), and appropriate anti-rabbit or anti-mouse HRP-linked secondary antibodies (1:500, 7074 and 7076, Cell Signaling). Blots were visualized with the iBright Imaging System (Thermo Fisher Scientific), and densitometry analysis was performed using Fiji ImageJ.

### RNA sequencing

RNA was isolated using the RNeasy Plus Mini Kit (Qiagen) according to the manufacturer’s protocol. Libraries were prepared and sequenced by the Center for Genome Technology, John P. Hussman Institute for Human Genomics, University of Miami Miller School of Medicine. Briefly, total RNA was prepared with the NuGen Universal Plus mRNA-Seq Kit (M01442 v2) using 50 ng of RNA, measured by Qubit, and amplified with 17 PCR cycles. Libraries were sequenced on an Illumina NovaSeq 6000, generating >40M single-end 100-bp reads per sample. Next-generation sequencing quality was assessed using FastQC (v0.11.3). Reads were trimmed with Trim Galore or Trimmomatic, aligned to the human genome (hg38/GRCH38) using Hisat2 or STAR, and gene counts were generated using StringTie or RSEM. Differential expression analysis was performed with EdgeR, applying a false discovery rate (FDR) < 0.05 as the cutoff.

### Animal Model

Mice deficient in HIF-1α (B6.129-*Hif1a^tm3Rsjo^*/J) and Myh6-MCM (A1cf^Tg(Myh6-cre/Esr1*)1Jmk^) mice in which Cre expression is driven by the murine αMyosin Heavy Chain (αMHC) promoter exclusively in CMs in a tamoxifen-inducible manner were purchased from The Jackson Lab (Bar Harbour, ME), and colonies were established in-house. After a series of matings, mice homozygous for B6.129-*Hif1a^tm3Rsjo^*/J, and either hemizygous for Myh6-MCM (HIF-1α^fl/fl^;Cre/+) or WT (no Myh6-MCM; HIF-1α^fl/fl^;+/+) were obtained, and we performed comprehensive cardiac phenotyping and molecular assays on these mice to examine the role of HIF-1α on GHRH therapy response in HFpEF. C57/Bl6N mice were obtained from Charles River Lab and used as WT controls.

#### Tamoxifen/Peanut Oil Treatment

Eight to twelve-week-old male and female HIF-1α^fl/fl^; Cre/+ mice were administered 100 µl intraperitoneal injections of 20 mg/ml tamoxifen (Sigma-Aldrich) dissolved in peanut oil for 5 consecutive days to activate the Cre gene and remove the floxed sequence, to generate mice with *HIF-1α* null alleles exclusively in CMs (HIF-1α^CM^KO). Some HIF-1α^fl/fl^;+/+ were similarly treated to control for the effect of tamoxifen, and some HIF-1α^fl/fl^; Cre/+ and HIF-1α^fl/fl^;+/+ were administered 100 µl peanut oil for 5 consecutive days to control for the vehicle. These mice are referred to as WT mice since the floxed HIF-1α gene remains intact.

All animals treated with tamoxifen/peanut oil were allowed to recover for 2 weeks before baseline echocardiography was conducted to identify any potential cardiac dysfunction induced by the Cre expression or treatment. To confirm the cardiac-specific HIF-1α deletion, we used immunofluorescence staining of the myocardium from WT and HIF-1α^CM^KO (tamoxifen-treated) mice.

#### Myocardial Infarction

WT and HIF-1α^CM^ KO mice were subjected to myocardial infarction as previously described with minor modifications^21^. Briefly, anesthesia was induced by 3-5% isoflurane, mice were intubated, and anesthesia was maintained around 2% during the surgery. A small incision was made on the left side of the thorax to expose the heart, the pericardium was removed, and the left coronary descending artery was ligated using a Prolene 7.0 suture. The chest was closed in layers using an absorbable suture, and the animal was allowed to recover in a warm recovery cage before being returned to its own cage. Buprenorphine extended release (3.25 mg/kg) was given about 10 minutes prior to surgery and repeated if needed after 72 hours. Hearts were harvested and prepared for immunofluorescence staining to confirm the absence of HIF-1α expression in the cardiomyocytes.

#### Cardiometabolic HFpEF model

Two weeks after tamoxifen/peanut oil treatment, baseline echocardiography was performed, then a high-fat diet (HFD, D12492, Research Diets) and *N*ω-nitrol-arginine methyl ester (L-NAME, Cayman Chemical, 0.5-1 g/L in drinking water) *ad libitum* was initiated for 5 weeks to induce the two-hit cardiometabolic HFpEF model as described^22, 23^. Both HFD and L-NAME were replaced every 3 days. The HFpEF phenotype was confirmed at 5 weeks after the diet regimen started, as previously described^13, 22, 24–27^. The control group received normal chow and regular water.

#### Growth hormone-releasing hormone agonist (GHRH-agonist)/Vehicle Treatment

Animals continued the HFD+L-NAME regimen for an additional 4 weeks while also receiving daily subcutaneous injections of either vehicle or the GHRH-agonist, MR-356, which was synthesized by one of us^28^. The compound was suspended in 0.1% dimethyl sulfoxide (DMSO) and 10% propylene glycol and passed through a 0.2 µm calcium acetate filter. After 5 weeks of the HFD+L-NAME regimen, animals were randomized to receive daily subcutaneous injections of equal volumes (100 ml) of GHRH-agonist (MR-356, 100 µg/kg) or vehicle (0.1% DMSO in 10% propylene glycol in saline) during a 4-week period, as previously described^13^.

#### Echocardiography

A comprehensive characterization of cardiometabolic HFpEF was conducted, as we recently described^12, 13, 29^, using conventional echocardiography, tissue Doppler imaging (TDI) and speckle-tracking echocardiography (STE). Serial cardiac images were acquired at baseline and at weeks 5 and 9 using a Vevo 2100 (VisualSonics Inc.) high-resolution (18-38 MHz) ultrasound scanner (MS-400) to assess systolic and diastolic functions. Briefly, the animals were anesthetized with isoflurane (3-5%, induction chamber) and maintained with 0.5-2% isoflurane to achieve consistent and comparable heart rates during image acquisition. Physiological parameters, including heart rate, respiratory rate and core body temperature, were continuously monitored by a built-in monitoring system.

Two-dimensional (B-mode) and M-mode images from the parasternal long-axis and short-axis (at the papillary muscle level) and apical views were acquired during the exam for cardiac morphology and function. All acquired images were stored and further analyzed offline using Vevo LAB software version 5.6.1 (FUJIFILM, VisualSonics). LV end-diastolic (LVEDD) and end-systolic (LVESD) dimensions, end-diastolic (EDV) and end-systolic (ESV) volumes, posterior (PWT) and anterior wall thickness (AWT) were determined. Relative wall thickness in diastole (RWTd) was calculated by the following formula: RWTd=2*PWd/LVEDD. Cardiac structure and function were assessed from cine-images in the parasternal long-axis and short-axis views using an automated analysis system (AutoLV software, FUJIFILM, VisualSonics). Pulsed wave Doppler of mitral blood (E and A velocities, ejection time (ET) of early filling of mitral inflow, isovolumetric relaxation time (IVRT) and isovolumetric contraction time (IVCT), and tissue Doppler (E’ and A’ waves) velocity traces of mitral valve annulus at the septal level were obtained from the four-chamber view to assess diastolic function. The E/A and E/E’ ratios and myocardial performance index (MPI) were then calculated. For STE, three consecutive cardiac cycles were selected for analysis, and semi-automated tracing of the endocardial and epicardial border were obtained to determine global longitudinal strain (GLS).

#### Blood pressure (BP) measurements

Blood pressure was measured using a non-invasive tail-cuff system (BP-2000, Visitech Systems, Apex, NC) at baseline, at 5 weeks once HFpEF was established, and after 4 weeks of daily GHRH-agonist or placebo treatment, as we previously reported^13^.

#### Intra-peritoneal Glucose tolerance test (ipGTT)

Prior to the ipGTT, the mice were fasted for 5-6 hours (morning fast), as we previously described^13^. The test was performed using a small drop of blood (< 5 µl) obtained from a tail tip and placed on the test strip of the blood glucose monitor (AlphaTrack2, Zoetis). After obtaining the baseline glucose level (t=0), mice received glucose via intraperitoneal injection (2 g of dextrose/kg in saline). The blood glucose levels were measured 15, 30, 45, 60 and 120 minutes after glucose administration. At the end of the experiment, mice were placed in a clean cage with water and food *ad libitum* for recovery. Mice were monitored carefully to observe any abnormal behavior.

#### Exercise training

For physical training, mice were acclimated to treadmill exercise for at least 3 days before the actual experiments; and an exhaustion test was performed in the experimental groups of mice as previously reported^13, 30^ following the American Physiological Society’s Resource Book for the Design of Animal Exercise Protocols^31^, with minor modifications. Briefly, on the first day of training, mice ran on the treadmill (Columbus Instruments) starting at a warm-up speed of 1.5 to 3.0 m/min, then the treadmill speed was slowly increased to 8 m/min for ∼5 min; after which speed was increased to 9 m/min at 5 min, 10 m/min at 7 min, and the treadmill was stopped at 10 min. On the second and third days of training, mice ran starting at a warm-up speed of 10 m/min; the treadmill speed was increased to 11 m/min at 5 min, 12 m/min at 10 min, and stopped at 15 min. Before proceeding to the exercise exhaustion test, mice were allowed at least one full day with no exposure to the treadmill. All measurements were done at approximately the same time each day.

#### Exercise exhaustion test (EET)

Briefly, mice were run on the treadmill starting at a warm-up speed of 5 m/min for 4 min, after which the speed was increased to 14 m/min for 2 min. Speed was increased by 2 m/min every 2 min until the mouse was exhausted, i.e., unable to return to running within 10 sec of direct contact with an electric-stimulus grid. Running time was measured, and distance was calculated.

#### Hemodynamic evaluation (Pressure-volume loops)

Pressure-volume (PV) measurements were acquired using a 1.4F microtip pressure-volume catheter (PVR, Millar Instruments), as we previously described^13, 32^. Briefly, mice were induced and maintained with isoflurane, and body temperature was controlled (∼37°C) throughout the procedure. The left internal jugular vein was exposed and cannulated with a 30-gauge needle to provide fluid support. The right carotid artery was exposed to permit the catheter to advance into the LV. PV loops were recorded during steady-state and temporary inferior vena cava occlusion. The ventilator was stopped momentarily (∼10 s) to avoid breathing interference during measurements. The volumes were calibrated using volumes derived from echocardiographic measurements. Load-dependent and load-independent parameters were evaluated to provide detailed insights into cardiac performance. Data analysis was done using LabChart Pro version 8.130 software (ADInstruments).

#### Post-mortem analysis

After hemodynamic studies, blood was withdrawn, the animals (under deep anesthesia) were humanely euthanized, and their tissues were harvested for further analysis. The whole heart and lungs were weighed and normalized to tibial length. To determine the dry lung weight, the samples were kept in an oven at 37⁰C for at least 72 hours. Lung water content (LWC) was calculated by subtracting dry lung weight from the wet lung weight.

#### Immunostaining

The paraffin-embedded sections of whole mouse hearts were stained for HIF-1a (Cell Signaling, CS) to confirm the absence of HIF-1α expression in the cardiomyocytes. Briefly, heart sections were deparaffinized with xylene and rehydrated in alcohol series and water as we previously described^8, 12, 13, 21^. Nuclei were counter-stained with 4’,6-diamidino-2-phenylindole (DAPI, Invitrogen, Carlsbad, CA). Image analyses were completed using a fluorescent microscope (Olympus IX81, Olympus America Inc., Center Valley, PA) or a LSM710 Zeiss confocal laser-scanning module (Carl Zeiss MicroImaging, Germany).

#### Statistical analysis

Statistical analyses and data visualizations were performed using SAS version 9.4 (SAS Institute, Cary, NC, USA) and GraphPad Prism version 10.4.0 (GraphPad Software, Boston, MA, USA). Categorical variables were presented as frequencies (percentage) and analyzed using Fisher’s exact test to compare between groups. Continuous variables were presented as a mean ± standard deviation or median [interquartile range], depending on their distribution (via Shapiro-Wilk normality test). Data was analyzed using Student’s T-Test and One and Two Way ANOVA followed by Tukey post hoc test, or Mann-Whitney U Test and Kruskal-Wallis test with Dunn’s multiple comparisons test if the normality test failed. For serial echocardiographic measurements, repeated measures ANOVA was used. A p-value < 0.05 was considered statistically significant.

## Results

### Transcriptomic profiling reveals activation of HIF1A signaling as a key downstream target of GHRH

We and others have shown that GHRHR activation has cardioprotective effects in animal models of both HFrEF and HFpEF ^8, 9, 11–13^. We sought to identify underlying molecular pathways responsible for the salutary effects of GHRH-R activation in the cardiac myocyte. To identify candidate signaling mechanism(s) specific to cardiomyocytes underlying these protective effects, we performed RNA-seq after short-term stimulation of human iPSC-derived cardiomyocytes (hiPSC-CM) with GHRH. We stimulated hiPSC-CMs with increasing doses (0 μM (control), 0.15 μM and 0.3 μM) of recombinant human GHRH (rhGHRH) for 45 min to identify immediate-early gene expression changes. GHRHR stimulation identified dose-dependent transcriptional changes in 482 genes **(Fig 1A)**. KEGG pathway enrichment analysis revealed increased expression of genes regulated by the ATF-2 (Activating Transcription Factor 2), HIF1A (Hypoxia-Inducible Factor 1-alpha), and HIF1B signaling pathway. ATF-2 and HIF1 genes are transcription factors that play important roles in cellular responses to hypoxia, with HIF1A being the most studied, considered more involved in the initial, acute hypoxic response. Of note, ATF-2 works to enhance the stability of HIF1A’s stability and transcriptional activity, and binds to the same DNA sequences cooperating with to HIF1α regulation of gene expression in response to hypoxia^33^. To further validate this pathway, we found HIF1A was significantly up-regulated in the RNA-seq data **(Fig. 1B)**. Orthogonal validation of these results by qPCR and immunoblotting validated the increased HIF1A RNA-(20-fold, **Fig. 1C**) and protein levels (3-4-fold, **Fig. 1D-E**) following GHRH stimulation. Given the prominent role of HIF1A in cardiac protection during hypoxic events, these results highlight a potential novel clinically targetable pathway,

**Figure 1.**
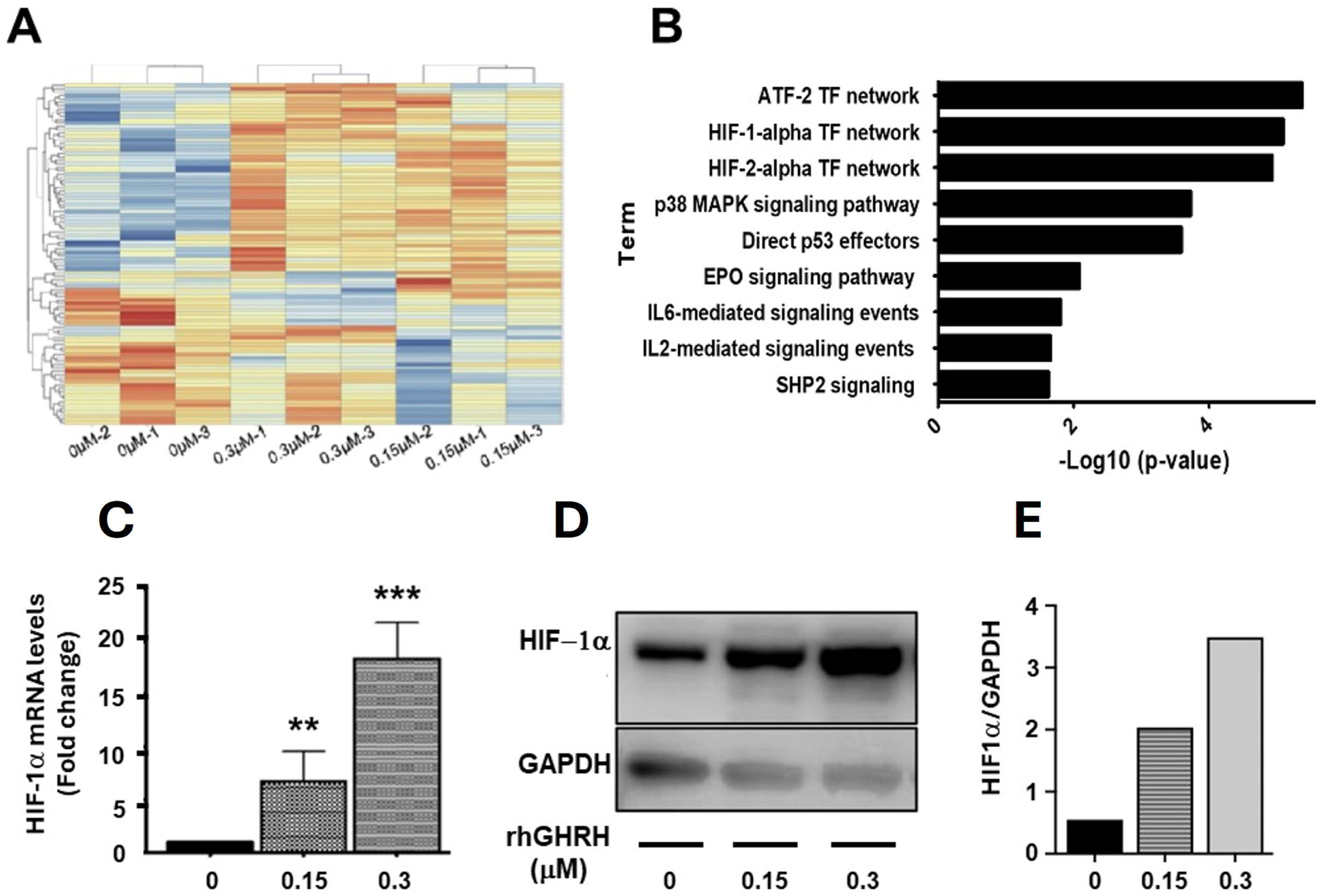
Changes in gene expression in iPSC-derived cardiomyocytes following short-term (45 min) exposure to GHRH. **(A)** Unsupervised hierarchical clustering heatmap of RNA-seq datasets. **(B)** Top 10 pathways and transcription factor (TF) networks of differentially expressed genes (DEGs). Fold change in HIF-1α RNA **(C)** and protein **(D, E)** expression in response to increasing concentrations of rhGHRH in a hypoxia independent manner. **p< 0.01, ***p<0.001 compared to 0 µM. N=3 in duplicate.

### Generation of inducible and cardiomyocyte-specific HIF-1α^CM^ KO mice

We next sought to test the hypothesis that HIF-1α signaling is required for GHRH-agonist-mediated reversal of the cardiometabolic HFpEF phenotype in vivo and generated a murine model of inducible cardiomyocyte-selective HIF-1α knockout (HIF-1α^CM^KO **(Fig. 2A)**, by crossing a HIF-1α^lox/flox^ with a Myh6-Mer-Cre-Mer^+/+^ strain. HIF-1a is expressed at high levels as a protective response to cardiac hypoxia/ischemia^34^ in various cardiac cell types. To investigate the expression of HIF-1α under normal and hypoxic conditions in our models, WT and HIF-1α^CM^KO mice were subjected to myocardial infarction (MI) by permanent ligation of the left anterior coronary artery. Infarcted hearts from WT mice (positive control, n=2) showed increased expression of HIF-1α in both cardiomyocytes (CMs) and non-CMs, including interstitial and vascular cells. In contrast, in HIF-1α^CM^KO mice (n=3), HIF-1α expression was detected only in non-cardiomyocytes **(Fig. 2B)**.

**Figure 2.**
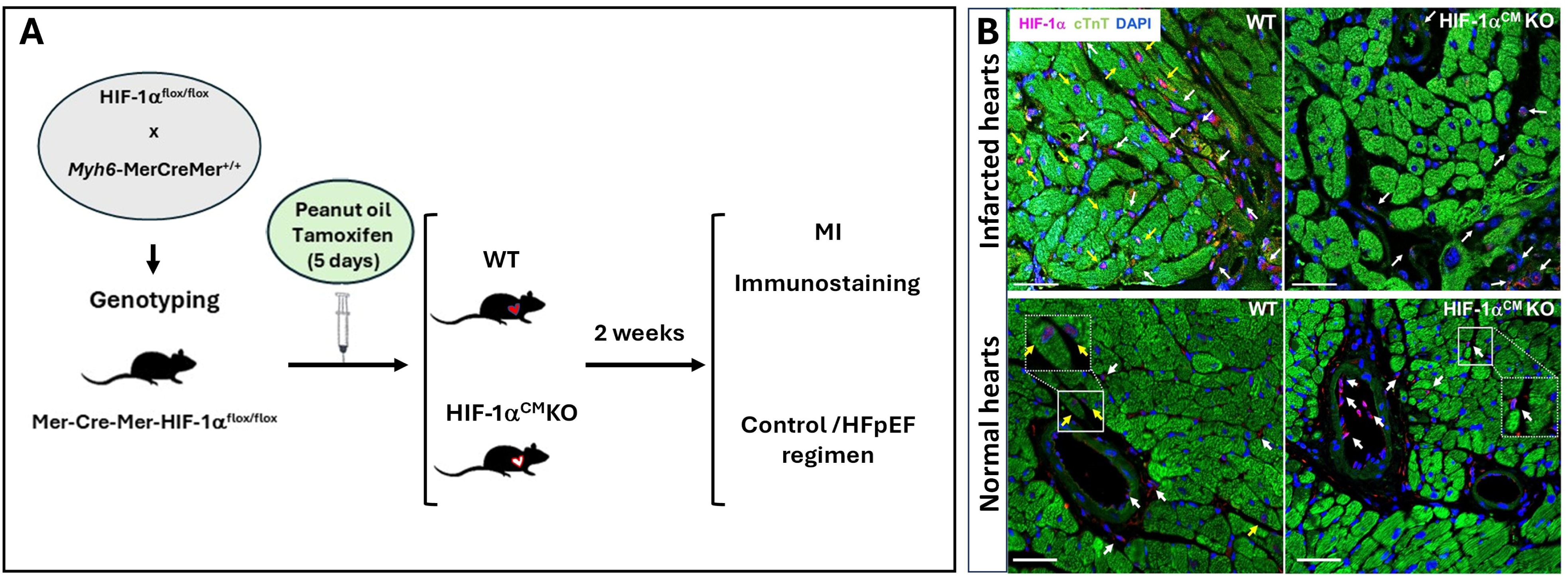
Generation of inducible and cardiomyocyte specific HIF-1α in HIF-1α^CM^ KO mice. **(A)** Schematic representation of cardiac-specific conditional and tamoxifen-inducible genetic deletion of HIF-1α in mice. **(B)** Representative immunofluorescence staining showing expression of HIF-1α (magenta), cardiac troponin T (cTnT, green) and nuclei (4′,6-diamidino-2-phenylindole: DAPI, blue). Top panels represent WT and HIF-1α^CM^KO infarcted hearts (n=2-3, respectively), while the bottom panels show normal hearts (n=2-3, respectively). HIF-1α is highly expressed in WT infarcted hearts (positive control). Yellow arrows indicate HIF-1α positive cardiomyocytes (CMs), and white arrows indicate non-cardiomyocytes (non-CMs, including interstitial and vascular cells). Insets at the bottom show higher magnification of HIF-1αpositive CMs in the WT heart (left panel) and non-CM in the HIF-1α^CM^KO heart (right panel), respectively. Scale bar: 50 mm.

### Cardiac phenotyping shows no baseline difference in cardiac function in cardiomyocyte-specific HIF-1α knockout mice (HIF-1α^CM^ KO)

HIF-1α knockout was induced in mice by injection of tamoxifen for 5 days, at least 2 weeks before HFpEF was induced using a HFD+L-NAME regimen for 5 weeks, as described previously^13, 22, 24^. Baseline characterization of cardiac function, including ejection fraction (EF), cardiac output (CO), relative wall thickness in diastole (RWTd), isovolumetric relaxation time (IVRT), myocardial performance index (MPI), and peak E/E’ ratio, revealed no significant differences between WT and HIF-1α^CM^ KO mice **(Fig. 3A-F)**, suggesting that our selected dose and frequency of tamoxifen were adequate to avoid cardiac toxicity. To reduce the potentially confounding effects of tamoxifen/αMHC MerCreMer, we initiated the HFD+L-NAME regimen two weeks after the last tamoxifen injection.

**Figure 3.**
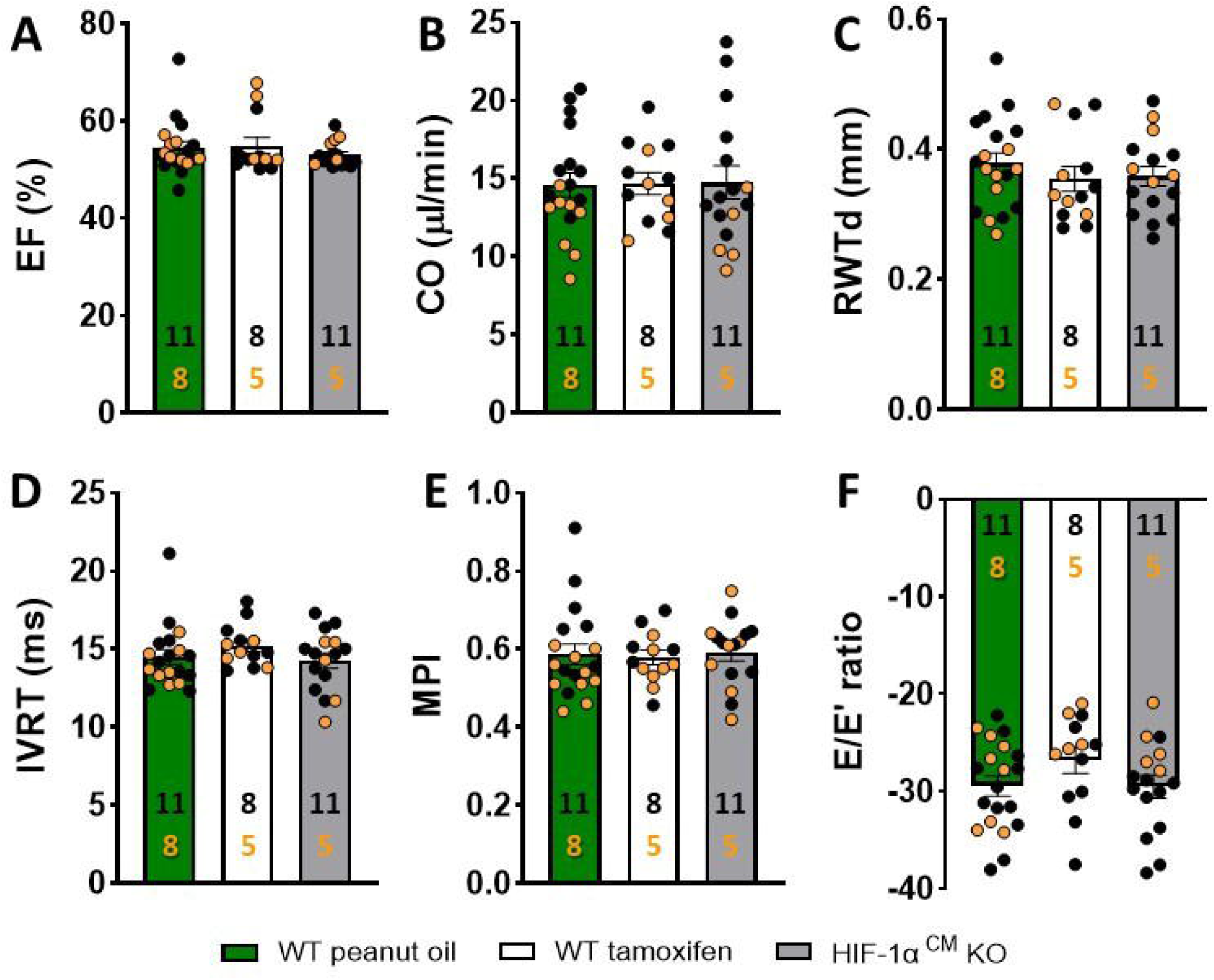
Cardiac phenotype of WT and HIF-1α^CM^KO mice at baseline. Echocardiographic evaluation revealed no effect of tamoxifen effect on cardiac performance in either WT or HIF-1α^CM^KO mice before the initiation of the control or HFD+L-NAME regime. Bar graphs correspond to **(A)** ejection fraction (EF), **(B)** cardiac output (CO), **(C)** relative wall thickness in diastole (RWTd), **(D)** isovolumetric relaxation time (IVRT), **(E)** myocardial performance index (MPI) and (F) peak E/E’ ratio. Female mice are represented by orange circles.

### WT and HIF-1α^CM^KO mice exhibit identical cardiometabolic HFpEF phenotype

Subsequently, the mice were randomly assigned to either the control or HFpEF diet, as shown in the experimental design **(Supplemental Fig. S1)**. Echocardiographic measurements revealed a prominent and characteristic HFpEF phenotype following 5 weeks of the HFD+L-NAME regimen. Both WT and HIF-1α^CM^KO mice exhibited hypertension, glucose intolerance, impaired diastolic function, and reduced exercise tolerance **(Fig. 4 A-H)**, whereas EF was preserved in both the control and HFpEF groups. Together, we observed no significant difference caused by cardiomyocyte specific HIF-1α loss on the development of the HFpEF phenotype.

**Figure 4.**
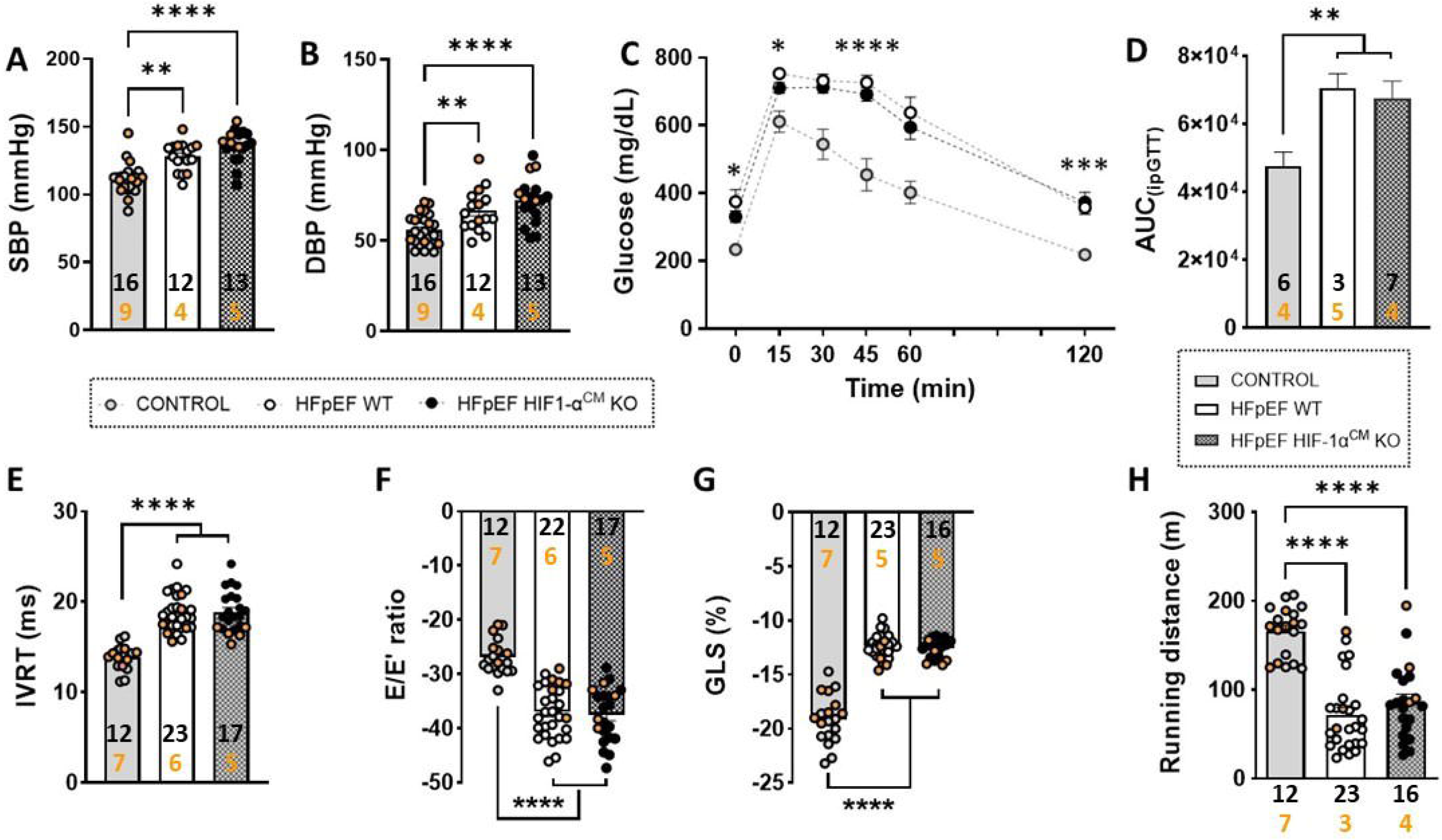
Systemic and cardiac changes at 5 weeks of HFD + L-NAME regimen. WT and HIF-1α^CM^ KO mice exhibit an identical cardiometabolic HFpEF phenotype, demonstrated by similar increases in blood pressure (**A-B**, **p<0.01 and ****p<0.0001 and glucose levels (**C:** Two-Way Repeated Measures (RM) ANOVA followed by Tukey’s test: *p<0.05, ***p<0.001 and ****p<0.0001, **D**: Unpaired t test, **p<0.01; elevated isovolumetric relaxation time (IVRT, **E**) and peak E wave velocity and peak E’ velocity (E/E’, **F**) ratio. Additionally, there was a severe decline in global longitudinal strain (GLS, **G**) and running capacity **(H)** compared to control mice. Female mice are represented by orange circles. (**E, F and G:** One-Way ANOVA followed by Tukey’s test and **H:** Kruskal-Wallis followed by Dunn’s test: ****p<0.0001.

### Cardiomyocyte-specific deletion of HIF-1α blocks the cardioprotective effects of GHRH-agonist in cardiometabolic HFpEF model

Next, we treated HFpEF-induced mice with either the GHRH agonist, MR-356, or placebo for 4 weeks. In agreement with our previous study, both systolic blood pressure **(Fig. 5A),** and glucose levels **(Fig. 5B-C)** were increased in all HFpEF groups and did not improve with GHRH-A treatment compared to the control group. In contrast, four weeks of treatment with the GHRH agonist MR-356 markedly improved key features of the cardiometabolic HFpEF phenotype in WT mice. The increase in isovolumetric relaxation time (IVRT, **Fig. 6A**) was attenuated in WT mice treated with MR-356, while the elevated E/E’ ratio **(Fig. 6B)** was restored to control levels. Similarly, global longitudinal strain (GLS, **Fig. 6C**) evaluated by speckle-strain echocardiography (STE) showed a severe decline in HFpEF mice. MR-356 markedly improved GLS in WT mice; but not in the HIF-1α^CM^KO mice, suggesting that HIF-1α is required for the therapeutic effects of GHRH-agonist, MR-356. In accordance with previous studies^13, 22, 23^, HFpEF mice exhibited a significant decrease in running distance during a treadmill exercise exhaustion test (EET) compared to controls. Notably, treatment with the GHRH agonist, MR-356, effectively improved the exercise capacity exclusively in WT mice **(Fig. 6D**), consistent with our previous results^13^. Although WT MR-356 mice showed higher exercise capacity than any other HFpEF group, it was not significantly different from HIF-1α^CM^KO mice treated with placebo or MR-356.

**Figure 5.**
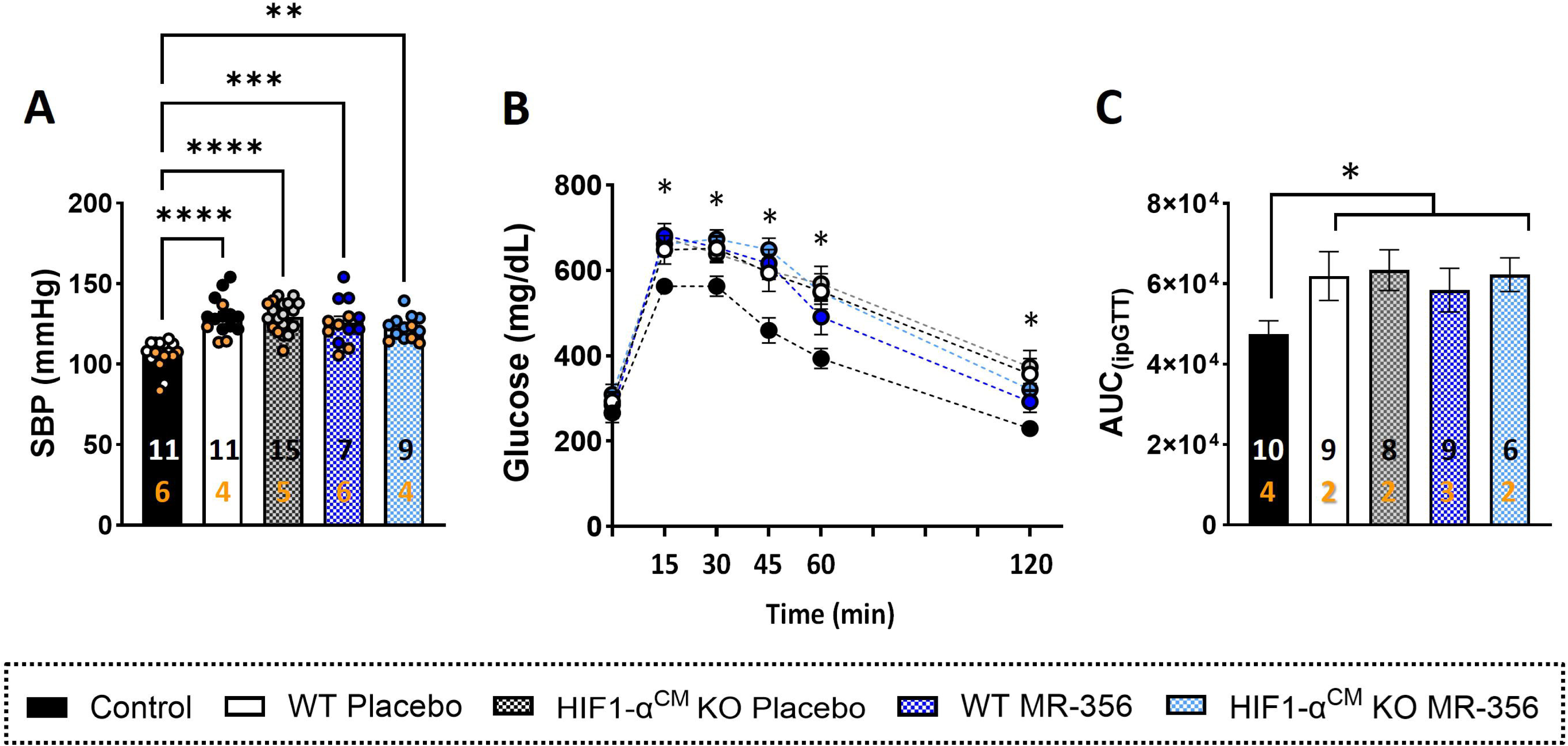
Effect of cardiac-specific HIF-1α deletion on blood pressure and glucose metabolism. WT and HIF-1α^CM^KO HFpEF mice show persistently elevated systolic blood pressure (SBP, **A**) and intraperitoneal glucose tolerance test (ipGTT) results revealed increased glucose levels in HFpEF mice compared to control mice **(B)**. Accordingly, the area under the curve (AUC) of ipGTT **(C)** was elevated in HFpEF-groups, suggesting impaired glucose tolerance in both GHRH-agonist and placebo-treated groups after 4 weeks of treatment. Female mice are represented by orange circles.

**Figure 6.**
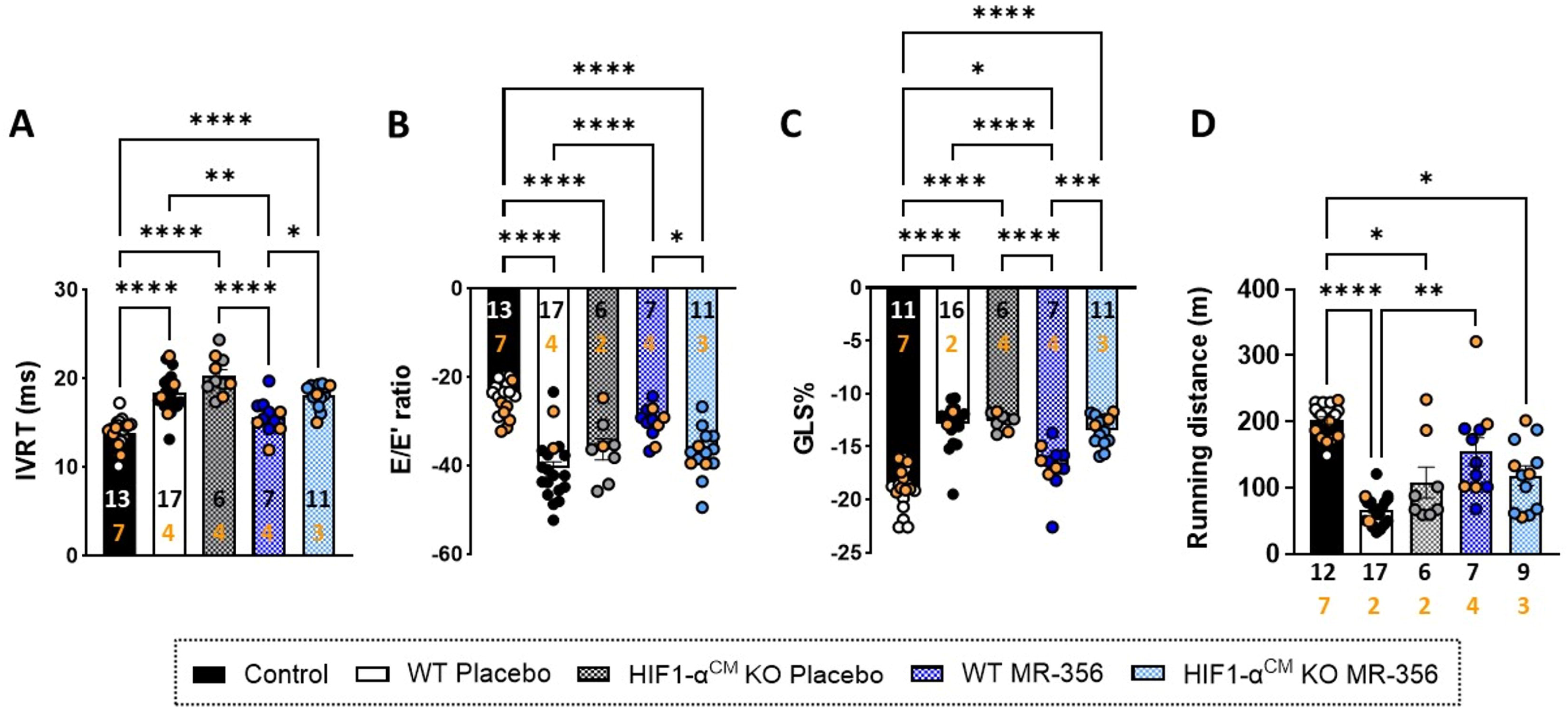
Cardiomyocyte-specific HIF-1α deletion blocks cardioprotective effects of GHRH-agonist, MR-356. Cardiometabolic HFpEF phenotype was attenuated by GHRH-agonist treatment in WT but not HIF-1α^CM^KO mice. Bar graphs represent: **(A)** Isovolumetric relaxation time (IVRT), (B) peak E/E’ ratio, **(C)** global longitudinal strain (GLS) and **(D)** exercise tolerance. The cardiometabolic HFpEF phenotype was attenuated or reversed by GHRH-agonist, MR-356 treatment in WT but not HIF-1α^CM^ KO mice. Running capacity was substantially reduced in HFpEF mice and greatly improved by GHRH-agonist, MR-356 in WT mice and it was mildly higher than both HIF-1α^CM^KO mice groups. Female mice are represented by orange circles. (**A-C**: One-Way ANOVA followed by Tukey’s test and **D:** Kruskal-Wallis followed by Dunn’s test, *p<0.05, **p<0.01, **p<0.001 and ****p<0.0001).

Pressure-volume (PV) analysis **(Fig. 7 and Supplemental Table S1)** revealed that end-diastolic pressure (EDP, **Fig. 7A**) and the slope of the end-diastolic pressure-volume relationship (EDPVR, **Fig. 7B**) were substantially higher in HFpEF mice than in those receiving the control diet, in agreement with diastolic dysfunction observed by echocardiography. MR-356 treatment significantly improved diastolic function in WT HFpEF mice but not HIF-1α^CM^KO mice, indicating that MR-356 treatment reduced ventricular stiffness **(Fig. 7 A-F)**.

**Figure 7.**
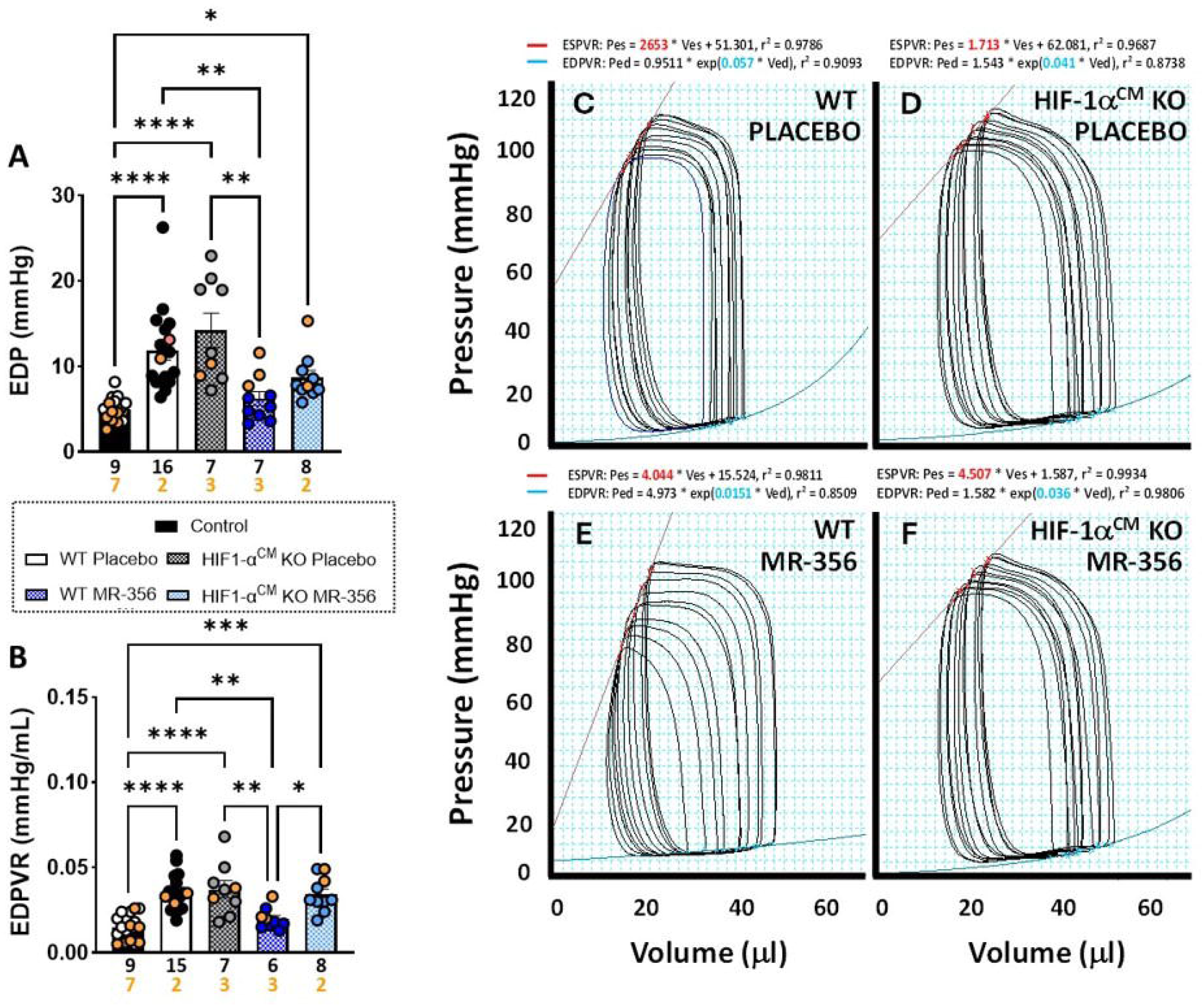
Hemodynamic changes in HFpEF mice after treatment with GHRH-agonist, MR-356. Left ventricle end-diastolic pressure (EDP, **A**) and the slope of the end-diastolic pressure-volume relationship (EDPVR, **B**) were markedly increased in HFpEF mice and restored to control levels in the WT MR-356 treated group, suggesting a remarkable improvement on diastolic function (**A**: Kruskall-Wallis followed by Dunn’s test **p<0.01 vs. both WT and HIF-1α^CM^KO Placebo mice; **B**: One-Way ANOVA followed by Tukey’s test *p<0.05, **p<0.01 vs. HIF-1α^CM^KO MR-356 and WT and HIF-1α^CM^KO Placebo mice, respectively). Representative pressure-volume (PV) loops show the linear regression for end-systolic pressure-volume relationship (ESPVR, red line) and exponential end-diastolic pressure-volume relationship (EDPVR, blue line) in male mice fed HFD+L-NAME. WT HFpEF mice **(C)** treated with **(D)** placebo exhibit an elevated slope of EDPVR (0.057 mmHg/µl), whereas **(E)** the GHRH-agonist, MR-356 restores the EDPVR toward normal (0.015 mmHg/µl). In contrast, in mice with a cardiac-specific deletion of HIF-1α (HIF-1α^CM^KO; **D and F**), administration of **(D)** placebo or **(F)** MR-356 fails to attenuate the EDPVR (0.04 mmHg/µl and 0.036 mmHg/µl, respectively).

Lastly, gravimetric measurements such as body weight (BW), heart weight (HW), lung weight (LW), heart weigh/tibia length ratio (HW/TL), lung weight/tibia length ratio (LW/TL), and lung water content (LWC) were assessed to evaluate cardiac hypertrophy and lung congestion **(Supplemental Table S2)**. As expected, BW substantially increased in all mice receiving the HFD+L-NAME regimen. These mice developed cardiac hypertrophy and pulmonary congestion demonstrated by increases in HW/TL, LW, LW/TL, and LWC **(Supplemental Figure S2)**. Importantly, treatment with GHRH-agonist MR356 reduced cardiac hypertrophy and ameliorate lung congestion in WT but not HIF-1α^CM^KO mice.

## Discussion

HFpEF remains a major clinical challenge due to its complex, multifactorial pathophysiology and lack of effective targeted therapies^1, 2^. In this study, we mechanistically delineated a novel cardioprotective pathway wherein activation of cardiac GHRHR directly induces HIF-1α signaling in cardiomyocytes under normoxic conditions. This oxygen-independent upregulation of HIF-1α represents a fundamental advance in understanding myocardial adaptation in HFpEF, as HIF-1α has been predominantly recognized for its role in hypoxic stress responses.

Through integrative transcriptomic and functional analyses, we showed that GHRHR activation induces HIF-1α expression in hiPSC-dervied cardiomyocytes, driving a cardioprotective transcriptional program. Functionally, this GHRHR-HIF-1α axis promotes metabolic reprogramming toward glycolysis, attenuated diastolic dysfunction, and improved myocardial deformation and exercise capacity. The importance of HIF-1α for the therapeutic effects of GHRHR agonists was confirmed using a cardiomyocyte-specific HIF-1α knockout model, which completely abolished the beneficial effects of GHRHR agonist treatment, including the improvements in diastolic parameters, global longitudinal strain, exercise tolerance, and cardiac remodeling.

Our findings position the GHRHR–HIF-1α axis as a central regulatory pathway integrating metabolic and contractile remodeling in HFpEF. Moreover, the ability to activate HIF-1α signaling independently of hypoxia suggests new opportunities to therapeutically reprogram cardiomyocyte metabolism and function in HFpEF patients, potentially synergizing with existing treatments such as SGLT2 inhibitors. Collectively, our mechanistic insights provide a strong foundation for the translational development of GHRHR agonists as treatment options for HFpEF. Here, we identify increased HIF1A signaling as a clinically targetable pathway, that can be activated in cardiomyocytes even under normoxic conditions through GRHR. We find prominent ATF and HIF1A gene signatures following short-term stimulation of hiPSC-CM with GHRHR. While the protective role of HIF-1α is well-validated in hypoxia, cardiac metabolism^35^, angiogenesis^36^, and diastolic distress^37^, less is known for ATF2. However, its known role in stabilizing HIF1A highlights the significant HIF1A response following GHRHR activation^33^ highlights a string HIF1A-mediated response.^38^ GHRH may activate ATF2 through cAMP/MAPK signaling, contributing to the upregulation of HIF-1α and its downstream effects.^33, 39^

HIF-1α is a key regulator of cardiac protection during hypoxia/ischemia^34^, modulating numerous metabolic pathways and energy generation processes.^40^ Its expression increases in ischemic heart disease^41^ and activates transcriptional programs that promote angiogenesis, metabolic reprogramming and cell survival. In addition to hypoxia, HIF-1α can be upregulated by growth factor signaling pathways, such as the PI3K/Akt/mTOR axis.^40^

Our data demonstrate that GHRH-agonist–induced upregulation of HIF-1α in cardiomyocytes not only restores diastolic function (as evidenced by normalization of IVRT and E/E′; Figs. 6A–B) but also improved myocardial deformation (GLS; Fig. 6C) and exercise capacity (Fig. 6D). These effects likely reflect the well-established role of HIF-1α in reprogramming substrate utilization toward more oxygen-efficient glycolysis and enhancing expression of key calcium-handling proteins (e.g., SERCA2a and phospholamban), thereby optimizing excitation– contraction coupling under stress conditions. By linking GHRHR activation to an oxygen-independent HIF-1α axis, our findings extend prior work on GHRH in ischemic injury and position HIF-1α as a nodal convergence point for metabolic and contractile rescue in HFpEF.

Our findings are in agreement with clinical data from studies using left ventricular biopsies obtained from patients with HFpEF. In these analyses, transcriptomic and metabolic^42^ evaluations identified HIF-1α as a dysregulated regulatory node (regulome)^43, 44^, further reinforcing its central role in HFpEF pathophysiology. Further support for our conclusions arise from studies of the mechanism underlying SGLT2i effects. Currently, SGLT2 inhibitors represent a major advancement HFpEF treatment^45–47^, offering notable cardiovascular benefits. While their mechanisms are still being elucidated, emerging evidence suggests they may induce a hypoxia-like state and activate HIF-1α–associated pathways^48–53^, potentially improving cardiac energetics and function by restoring and maintaining cellular homeostasis and survival^53^. This supports the broader therapeutic relevance of targeting HIF-1α–related mechanisms in HFpEF.

The emergence of SGLT2 inhibitors as foundational therapy in HFpEF highlights the importance of targeting myocardial energy balance and cellular homeostasis^45, 54^. Mechanistic studies propose that SGLT2i induce a pseudohypoxic state^55–57^, culminating in HIF-1α stabilization and downstream angiogenic, anti-fibrotic, and antioxidant pathways. Our demonstration that direct GHRHR agonism robustly activates HIF-1α in cardiomyocytes, even in normoxia suggests a complementary or synergistic avenue to amplify these pathways. Combining GHRH-agonists with SGLT2i could potentially harness dual mechanisms—renal-mediated hemodynamic unloading and direct myocardial HIF-1α–dependent metabolic reprogramming—warranting dedicated preclinical trials to evaluate additive benefits. Additionally, our recent findings reveal that myocardial HIF-1α signaling is developmentally regulated via an oxygen-independent mechanism involving autocrine stimulation of the GHRH signaling pathway, specifically in NKX2-5-expressing cardiomyocytes^20^. These results align with our preclinical studies demonstrating that GHRH agonists reverse the HFpEF phenotype in two distinct murine^12, 13^ and a swine^14^ model of HFpEF, including the restoration of CM Ca^2+^ handling abnormalities^12^. Moreover, our preclinical studies have repeatedly shown that GHRH-agonists promote a comprehensive CM repair program in MI^8–12, 28^, and other forms of heart disease in both large and small animal models by protecting CMs from oxidative stress^28^ and enhancing the survival and proliferation of CMs and cardiac progenitor cells (CPCs) through activation of the Akt and Erk pathways.

These beneficial effects translate to improved cardiac function, increased peripheral vascular density, reduced remodeling, increased cell proliferation, and attenuated fibrosis in rat^8, 28^, swine^9^ models of ischemic cardiomyopathy. Furthermore, GHRH agonists have shown efficacy in inhibiting vascular calcification^58^and mitigating hypertrophic cardiomyopathy^59^.

HFpEF is a complex, multifactorial syndrome influenced by sex, age and cardiovascular risk factors^60^. It is more prevalent in women; however, previous studies have reported that HFD+L-NAME-induced HFpEF is less effectively induced in female^23^ than in male mice. While our study primarily used male mice, we incorporated a subset of females for comparison. We found no significant sex-based differences, except in body size and exercise capacity. While the female group was underrepresented in certain treatment groups, both male and female, intact mice were responsive to GHRH agonist treatment, supporting its broader applicability. Nonetheless, further research is needed to clarify sex-specific mechanisms and the influence of hormonal factors in HFpEF^2^ pathophysiology and treatment response^61^.

We also observed a significant improvement in exercise capacity in GHRHR agonist treated animals. While this underscores the cardiac benefits of GHRH agonist in intact WT hearts, we cannot rule out the possibility that improvements in exercise tolerance may be due, in part, to extracardiac mechanisms.

In summary, our findings suggest that the activation of GHRH receptor signaling is an effective therapeutic strategy for treating the cardiometabolic HFpEF phenotype. We demonstrate that the beneficial effects of GHRHR agonists are mediated through HIF-1α expression, specifically in cardiomyocytes. While our transgenic model cannot fully replicate the entire HFpEF phenotype, it provides valuable mechanistic insights into cardiac contributors to the disease and suggests a potential novel therapeutic approach, and points towards new directions for future research. Further exploration using both in vitro and in vivo models will provide deeper insights into these interactions and their potential therapeutic synergies. This novel mechanism holds promise for developing targeted therapies to better serve this growing patient population.

## Acknowledgments

This work was supported by Cardiovascular Health Retreat Team Science Funding Program (to RMKT, SK and JH), NIH grant RO1 HL10710 (to JMH) and by Medical Research Service of the Veterans Affairs Department and the University of Miami Miller School of Medicine and funding from the Wallace H. Coulter Foundation, SB26 MT66142E (to AVS).

## Author Contributions

RMKT, WB, KEH and JMH designed the study; RMKT, LMT, WB, ACBAW, KEH performed research; RC, WS and AVS contributed new reagents; RMKT, LMT, ACBAW, KEH, SK, RS and SK analyzed data; RMKT, WB, SK, LAS, SK and JMH wrote the paper.

## SUPPLEMENTAL INFORMATION

**Supplemental Figure S1. Experimental design.** WT and HIF-1α^CM^KO mice will be fed a high-fat diet plus L-NAME in drinking water; HFD+L-NAME) *ad libitum* for 5 weeks to induce the cardiometabolic HFpEF phenotype. The diet regimen was continued for the 4-week period during which mice were administered a daily subcutaneous injection of vehicle control (Placebo) or the GHRH-agonist, MR-356. Heart failure with preserved ejection fraction (HFpEF), wild-type (WT), cardiomyocyte-specific deletion of HIF-1α (HIF-1α^CM^KO), male (♂, M), female (♀, F), high-fat diet (HFD), *N*ω-nitrol-arginine methyl ester (L-NAME), Growth hormone-releasing hormone agonist (GHRH-agonist, MR-356).

**Supplemental Figure S2. Gravimetric data.** Morphometric data are normalized to tibia length (TL). Values are expressed as means ± standard error of mean (SEM). Bar graphs correspond to **(A and B)** body weight (BW) of males and females, respectively; **(C)** ratio of heart weight to tibia length (HW/TL), **(D)** ratio of lung weight to tibia length (LW/TL), and **(E)** lung water content (LWC). Female mice are represented by orange circles **(C-E)**.

**Supplemental Table S1. Hemodynamic parameters**.

**Supplemental Table S2. Gravimetric data.**

## References

1. Cannata A and McDonagh TA. Heart Failure with Preserved Ejection Fraction. N Engl J Med. 2025;392:173–184.

2. Radakrishnan A, Agrawal S, Singh N, Barbieri A, Shaw LJ, Gulati M and Lala A. Underpinnings of Heart Failure With Preserved Ejection Fraction in Women - From Prevention to Improving Function. A Co-publication With the American Journal of Preventive Cardiology and the Journal of Cardiac Failure. J Card Fail. 2025.

3. Borlaug BA, Sharma K, Shah SJ and Ho JE. Heart Failure With Preserved Ejection Fraction: JACC Scientific Statement. J Am Coll Cardiol. 2023;81:1810–1834.

4. Bozkurt B, Ahmad T, Alexander K, Baker WL, Bosak K, Breathett K, Carter S, Drazner MH, Dunlay SM, Fonarow GC, Greene SJ, Heidenreich P, Ho JE, Hsich E, Ibrahim NE, Jones LM, Khan SS, Khazanie P, Koelling T, Lee CS, Morris AA, Page RL, 2nd, Pandey A, Piano MR, Sandhu AT, Stehlik J, Stevenson LW, Teerlink J, Vest AR, Yancy C and Ziaeian B. HF STATS 2024: Heart Failure Epidemiology and Outcomes Statistics An Updated 2024 Report from the Heart Failure Society of America. J Card Fail. 2025;31:66–116.

5. Savarese G, Schiattarella GG, Lindberg F, Anker MS, Bayes-Genis A, Bäck M, Braunschweig F, Bucciarelli-Ducci C, Butler J, Cannata A, Capone F, Chioncel O, D’Elia E, González A, Filippatos G, Girerd N, Hulot JS, Lam CSP, Lund LH, Maack C, Moura B, Petrie MC, Piepoli M, Shehab A, Yilmaz MB, Seferovic P, Tocchetti CG, Rosano GMC and Metra M. Heart failure and obesity: Translational approaches and therapeutic perspectives. A scientific statement of the Heart Failure Association of the ESC. Eur J Heart Fail. 2025.

6. Dulce RA, Hatzistergos KE, Kanashiro-Takeuchi RM, Takeuchi LM, Balkan W and Hare JM. Growth hormone-releasing hormone signaling and manifestations within the cardiovascular system. Rev Endocr Metab Disord. 2025.

7. Granata R, Leone S, Zhang X, Gesmundo I, Steenblock C, Cai R, Sha W, Ghigo E, Hare JM, Bornstein SR and Schally AV. Growth hormone-releasing hormone and its analogues in health and disease. Nat Rev Endocrinol. 2024.

8. Kanashiro-Takeuchi RM, Tziomalos K, Takeuchi LM, Treuer AV, Lamirault G, Dulce R, Hurtado M, Song Y, Block NL, Rick F, Klukovits A, Hu Q, Varga JL, Schally AV and Hare JM. Cardioprotective effects of growth hormone-releasing hormone agonist after myocardial infarction. Proc Natl Acad Sci U S A. 2010;107:2604–9.

9. Bagno LL, Kanashiro-Takeuchi RM, Suncion VY, Golpanian S, Karantalis V, Wolf A, Wang B, Premer C, Balkan W, Rodriguez J, Valdes D, Rosado M, Block NL, Goldstein P, Morales A, Cai RZ, Sha W, Schally AV and Hare JM. Growth hormone-releasing hormone agonists reduce myocardial infarct scar in swine with subacute ischemic cardiomyopathy. J Am Heart Assoc. 2015;4.

10. Kanashiro-Takeuchi RM, Szalontay L, Schally AV, Takeuchi LM, Popovics P, Jaszberenyi M, Vidaurre I, Zarandi M, Cai RZ, Block NL, Hare JM and Rick FG. New therapeutic approach to heart failure due to myocardial infarction based on targeting growth hormone-releasing hormone receptor. Oncotarget. 2015;6:9728–39.

11. Kanashiro-Takeuchi RM, Takeuchi LM, Rick FG, Dulce R, Treuer AV, Florea V, Rodrigues CO, Paulino EC, Hatzistergos KE, Selem SM, Gonzalez DR, Block NL, Schally AV and Hare JM. Activation of growth hormone releasing hormone (GHRH) receptor stimulates cardiac reverse remodeling after myocardial infarction (MI). Proc Natl Acad Sci U S A. 2012;109:559–63.

12. Dulce RA, Kanashiro-Takeuchi RM, Takeuchi LM, Salerno AG, Wanschel A, Kulandavelu S, Balkan W, Zuttion M, Cai R, Schally AV and Hare JM. Synthetic growth hormone-releasing hormone agonist ameliorates the myocardial pathophysiology characteristic of heart failure with preserved ejection fraction. Cardiovasc Res. 2023;118:3586–3601.

13. Kanashiro-Takeuchi RM, Takeuchi LM, Dulce RA, Kazmierczak K, Balkan W, Cai R, Sha W, Schally AV and Hare JM. Efficacy of a growth hormone-releasing hormone agonist in a murine model of cardiometabolic heart failure with preserved ejection fraction. Am J Physiol Heart Circ Physiol. 2023;324:H739–h750.

14. Rieger AC, Bagno LL, Salerno A, Florea V, Rodriguez J, Rosado M, Turner D, Dulce RA, Takeuchi LM, Kanashiro-Takeuchi RM, Buchwald P, Wanschel A, Balkan W, Schulman IH, Schally AV and Hare JM. Growth hormone-releasing hormone agonists ameliorate chronic kidney disease-induced heart failure with preserved ejection fraction. Proc Natl Acad Sci U S A. 2021;118.

15. Semenza GL. Regulation of metabolism by hypoxia-inducible factor 1. Cold Spring Harb Symp Quant Biol. 2011;76:347–53.

16. Iyer NV, Kotch LE, Agani F, Leung SW, Laughner E, Wenger RH, Gassmann M, Gearhart JD, Lawler AM, Yu AY and Semenza GL. Cellular and developmental control of O2 homeostasis by hypoxia-inducible factor 1 alpha. Genes Dev. 1998;12:149–62.

17. Xia X, Lemieux ME, Li W, Carroll JS, Brown M, Liu XS and Kung AL. Integrative analysis of HIF binding and transactivation reveals its role in maintaining histone methylation homeostasis. Proc Natl Acad Sci U S A. 2009;106:4260–5.

18. Zheng J, Chen P, Zhong J, Cheng Y, Chen H, He Y and Chen C. HIF1alpha in myocardial ischemiareperfusion injury (Review). Mol Med Rep. 2021;23.

19. Guide for the Care and Use of Laboratory Animals 8th Edition; 2011.

20. Amarylis C.B.A. Wanschel; Konstantinos E. Hatzistergos; Alessandro G. Salerno JNK, Stefan Kurtenback, Daniel A. Rodriguez, Krystalenia Valasaki, Wayne Balkan, Derek Dykxhoorn, Andrew V. Schally, Joshua M. Hare. The Growth Hormone Releasing Hormone Signaling Pathway Governs Cardiomyocyte differentiation in human iPS cells. BioRxiv. 2022.

21. Hatzistergos KE, Paulino EC, Dulce RA, Takeuchi LM, Bellio MA, Kulandavelu S, Cao Y, Balkan W, Kanashiro-Takeuchi RM and Hare JM. S-Nitrosoglutathione Reductase Deficiency Enhances the Proliferative Expansion of Adult Heart Progenitors and Myocytes Post Myocardial Infarction. J Am Heart Assoc. 2015;4.

22. Schiattarella GG, Altamirano F, Tong D, French KM, Villalobos E, Kim SY, Luo X, Jiang N, May HI, Wang ZV, Hill TM, Mammen PPA, Huang J, Lee DI, Hahn VS, Sharma K, Kass DA, Lavandero S, Gillette TG and Hill JA. Nitrosative stress drives heart failure with preserved ejection fraction. Nature. 2019;568:351–356.

23. Tong D, Schiattarella GG, Jiang N, May HI, Lavandero S, Gillette TG and Hill JA. Female Sex Is Protective in a Preclinical Model of Heart Failure With Preserved Ejection Fraction. Circulation. 2019;140:1769–1771.

24. Schiattarella GG, Altamirano F, Kim SY, Tong D, Ferdous A, Piristine H, Dasgupta S, Wang X, French KM, Villalobos E, Spurgin SB, Waldman M, Jiang N, May HI, Hill TM, Luo Y, Yoo H, Zaha VG, Lavandero S, Gillette TG and Hill JA. Xbp1s-FoxO1 axis governs lipid accumulation and contractile performance in heart failure with preserved ejection fraction. Nat Commun. 2021;12:1684.

25. Tong D, Schiattarella GG, Jiang N, Altamirano F, Szweda PA, Elnwasany A, Lee DI, Yoo H, Kass DA, Szweda LI, Lavandero S, Verdin E, Gillette TG and Hill JA. NAD(+) Repletion Reverses Heart Failure With Preserved Ejection Fraction. Circ Res. 2021;128:1629–1641.

26. LaPenna KB, Li Z, Doiron JE, Sharp TE, 3rd, Xia H, Moles K, Koul K, Wang JS, Polhemus DJ, Goodchild TT, Patel RB, Shah SJ and Lefer DJ. Combination Sodium Nitrite and Hydralazine Therapy Attenuates Heart Failure With Preserved Ejection Fraction Severity in a “2-Hit” Murine Model. J Am Heart Assoc. 2023;12:e028480.

27. Smolgovsky S, Bayer AL, Kaur K, Sanders E, Aronovitz M, Filipp ME, Thorp EB, Schiattarella GG, Hill JA, Blanton RM, Cubillos-Ruiz JR and Alcaide P. Impaired T cell IRE1α/XBP1 signaling directs inflammation in experimental heart failure with preserved ejection fraction. J Clin Invest. 2023;133.

28. Cai R, Schally AV, Cui T, Szalontay L, Halmos G, Sha W, Kovacs M, Jaszberenyi M, He J, Rick FG, Popovics P, Kanashiro-Takeuchi R, Hare JM, Block NL and Zarandi M. Synthesis of new potent agonistic analogs of growth hormone-releasing hormone (GHRH) and evaluation of their endocrine and cardiac activities. Peptides. 2014;52:104–12.

29. Dulce RA, Kanashiro-Takeuchi RM, Takeuchi LM, Salerno AG, Wanschel A, Kulandavelu S, Balkan W, Zuttion M, Cai R, Schally AV and Hare JM. Synthetic growth hormone-releasing hormone agonist ameliorates the myocardial pathophysiology characteristic of HFpEF. Cardiovasc Res. 2022.

30. Colbert LH, Davis JM, Essig DA, Ghaffar A and Mayer EP. Tissue expression and plasma concentrations of TNFalpha, IL-1beta, and IL-6 following treadmill exercise in mice. Int J Sports Med. 2001;22:261–7.

31. Kregel Kadbffmhe, Musch, TI, O; Leary DS; Parks CM; Poole, DC; Ra’anan AW; Sheriff DD; Sturek MS; Toth, LA. Resource Book for the Design of Animal Exercise Protocols; 2006.

32. Williams M, Kamiar A, Condor Capcha JM, Rasmussen MA, Alitter Q, Kanashiro Takeuchi R, Mitsuru Takeuchi L, Hare JM and Shehadeh LA. A Murine Model of Hyperlipidemia-Induced Heart Failure with Preserved Ejection Fraction. J Vis Exp. 2024.

33. Choi JH, Cho HK, Choi YH and Cheong J. Activating transcription factor 2 increases transactivation and protein stability of hypoxia-inducible factor 1alpha in hepatocytes. Biochem J. 2009;424:285–96.

34. Cerychova R and Pavlinkova G. HIF-1, Metabolism, and Diabetes in the Embryonic and Adult Heart. Frontiers in endocrinology. 2018;9:460–460.

35. Semenza GL. Hypoxia-inducible factor 1 and cardiovascular disease. Annu Rev Physiol. 2014;76:39–56.

36. Bakleh MZ and Al Haj Zen A. The Distinct Role of HIF-1α and HIF-2α in Hypoxia and Angiogenesis. Cells. 2025;14.

37. Sato T and Takeda N. The roles of HIF-1α signaling in cardiovascular diseases. J Cardiol. 2023;81:202–208.

38. Bhoumik A and Ronai Z. ATF2: a transcription factor that elicits oncogenic or tumor suppressor activities. Cell Cycle. 2008;7:2341–5.

39. Bhoumik A, Takahashi S, Breitweiser W, Shiloh Y, Jones N and Ronai Z. ATM-dependent phosphorylation of ATF2 is required for the DNA damage response. Mol Cell. 2005;18:577–87.

40. Bishop T and Ratcliffe PJ. HIF hydroxylase pathways in cardiovascular physiology and medicine. Circulation research. 2015;117:65–79.

41. Stanley WC, Recchia FA and Lopaschuk GD. Myocardial substrate metabolism in the normal and failing heart. Physiological reviews. 2005;85:1093–1129.

42. Warbrick I and Rabkin SW. Hypoxia-inducible factor 1-alpha (HIF-1α) as a factor mediating the relationship between obesity and heart failure with preserved ejection fraction. Obes Rev. 2019;20:701–712.

43. Das S, Frisk C, Eriksson MJ, Walentinsson A, Corbascio M, Hage C, Kumar C, Asp M, Lundeberg J, Maret E, Persson H, Linde C and Persson B. Transcriptomics of cardiac biopsies reveals differences in patients with or without diagnostic parameters for heart failure with preserved ejection fraction. Sci Rep. 2019;9:3179.

44. Hahn VS, Petucci C, Kim MS, Bedi KC, Jr., Wang H, Mishra S, Koleini N, Yoo EJ, Margulies KB, Arany Z, Kelly DP, Kass DA and Sharma K. Myocardial Metabolomics of Human Heart Failure With Preserved Ejection Fraction. Circulation. 2023;147:1147–1161.

45. Anker SD, Butler J, Filippatos G, Ferreira JP, Bocchi E, Böhm M, Brunner-La Rocca HP, Choi DJ, Chopra V, Chuquiure-Valenzuela E, Giannetti N, Gomez-Mesa JE, Janssens S, Januzzi JL, Gonzalez-Juanatey JR, Merkely B, Nicholls SJ, Perrone SV, Piña IL, Ponikowski P, Senni M, Sim D, Spinar J, Squire I, Taddei S, Tsutsui H, Verma S, Vinereanu D, Zhang J, Carson P, Lam CSP, Marx N, Zeller C, Sattar N, Jamal W, Schnaidt S, Schnee JM, Brueckmann M, Pocock SJ, Zannad F and Packer M. Empagliflozin in Heart Failure with a Preserved Ejection Fraction. N Engl J Med. 2021;385:1451–1461.

46. Solomon SD, McMurray JJV, Claggett B, de Boer RA, DeMets D, Hernandez AF, Inzucchi SE, Kosiborod MN, Lam CSP, Martinez F, Shah SJ, Desai AS, Jhund PS, Belohlavek J, Chiang CE, Borleffs CJW, Comin-Colet J, Dobreanu D, Drozdz J, Fang JC, Alcocer-Gamba MA, Al Habeeb W, Han Y, Cabrera Honorio JW, Janssens SP, Katova T, Kitakaze M, Merkely B, O’Meara E, Saraiva JFK, Tereshchenko SN, Thierer J, Vaduganathan M, Vardeny O, Verma S, Pham VN, Wilderäng U, Zaozerska N, Bachus E, Lindholm D, Petersson M and Langkilde AM. Dapagliflozin in Heart Failure with Mildly Reduced or Preserved Ejection Fraction. N Engl J Med. 2022;387:1089–1098.

47. McDonagh TA, Metra M, Adamo M, Gardner RS, Baumbach A, Böhm M, Burri H, Butler J, Čelutkienė J, Chioncel O, Cleland JGF, Coats AJS, Crespo-Leiro MG, Farmakis D, Gilard M, Heymans S, Hoes AW, Jaarsma T, Jankowska EA, Lainscak M, Lam CSP, Lyon AR, McMurray JJV, Mebazaa A, Mindham R, Muneretto C, Francesco Piepoli M, Price S, Rosano GMC, Ruschitzka F and Kathrine Skibelund A. 2021 ESC Guidelines for the diagnosis and treatment of acute and chronic heart failure. Eur Heart J. 2021;42:3599–3726.

48. Faivre A and de Seigneux S. The role of hypoxia in chronic kidney disease: a nuanced perspective. Curr Opin Nephrol Hypertens. 2024;33:414–419.

49. Packer M. SGLT2 inhibitors: role in protective reprogramming of cardiac nutrient transport and metabolism. Nat Rev Cardiol. 2023;20:443–462.

50. Packer M. Foetal recapitulation of nutrient surplus signalling by O-GlcNAcylation and the failing heart. Eur J Heart Fail. 2023;25:1199–1212.

51. Packer M. Mutual Antagonism of Hypoxia-Inducible Factor Isoforms in Cardiac, Vascular, and Renal Disorders. JACC Basic Transl Sci. 2020;5:961–968.

52. Packer M. Fetal Reprogramming of Nutrient Surplus Signaling, O-GlcNAcylation, and the Evolution of CKD. J Am Soc Nephrol. 2023;34:1480–1491.

53. Packer M. Role of Deranged Energy Deprivation Signaling in the Pathogenesis of Cardiac and Renal Disease in States of Perceived Nutrient Overabundance. Circulation. 2020;141:2095–2105.

54. Cinti F, Laborante R, Cappannoli L, Morciano C, Gugliandolo S, Pontecorvi A, Burzotta F, Donniacuo M, Cappetta D, Patti G, Giaccari A and D’Amario D. The effects of SGLT2i on cardiac metabolism in patients with HFpEF: Fact or fiction? Cardiovasc Diabetol. 2025;24:208.

55. Lopaschuk GD and Verma S. Mechanisms of Cardiovascular Benefits of Sodium Glucose Co-Transporter 2 (SGLT2) Inhibitors: A State-of-the-Art Review. JACC Basic Transl Sci. 2020;5:632–644.

56. Yurista SR, Silljé HHW, Oberdorf-Maass SU, Schouten EM, Pavez Giani MG, Hillebrands JL, van Goor H, van Veldhuisen DJ, de Boer RA and Westenbrink BD. Sodium-glucose co-transporter 2 inhibition with empagliflozin improves cardiac function in non-diabetic rats with left ventricular dysfunction after myocardial infarction. Eur J Heart Fail. 2019;21:862–873.

57. Verma S and McMurray JJV. SGLT2 inhibitors and mechanisms of cardiovascular benefit: a state-of-the-art review. Diabetologia. 2018;61:2108–2117.

58. Shen J, Zhang N, Lin YN, Xiang P, Liu XB, Shan PF, Hu XY, Zhu W, Tang YL, Webster KA, Cai R, Schally AV, Wang J and Yu H. Regulation of Vascular Calcification by Growth Hormone-Releasing Hormone and Its Agonists. Circ Res. 2018;122:1395–1408.

59. Gesmundo I, Miragoli M, Carullo P, Trovato L, Larcher V, Di Pasquale E, Brancaccio M, Mazzola M, Villanova T, Sorge M, Taliano M, Gallo MP, Alloatti G, Penna C, Hare JM, Ghigo E, Schally AV, Condorelli G and Granata R. Growth hormone-releasing hormone attenuates cardiac hypertrophy and improves heart function in pressure overload-induced heart failure. Proc Natl Acad Sci U S A. 2017;114:12033–12038.

60. Gallet R, de Couto G, Simsolo E, Valle J, Sun B, Liu W, Tseliou E, Zile MR and Marbán E. Cardiosphere-derived cells reverse heart failure with preserved ejection fraction (HFpEF) in rats by decreasing fibrosis and inflammation. JACC Basic Transl Sci. 2016;1:14–28.

61. Sotomi Y, Hikoso S, Nakatani D, Mizuno H, Okada K, Dohi T, Kitamura T, Sunaga A, Kida H, Oeun B, Sato T, Komukai S, Tamaki S, Yano M, Hayashi T, Nakagawa A, Nakagawa Y, Yasumura Y, Yamada T and Sakata Y. Sex Differences in Heart Failure With Preserved Ejection Fraction. J Am Heart Assoc. 2021;10:e018574.

