## Supplementary figures and images for "Cardiomyocyte-specific expression of HIF-1α mediates the cardioprotective effects of Growth Hormone Releasing Hormone (GHRH)"

### Supplemental Figure S1

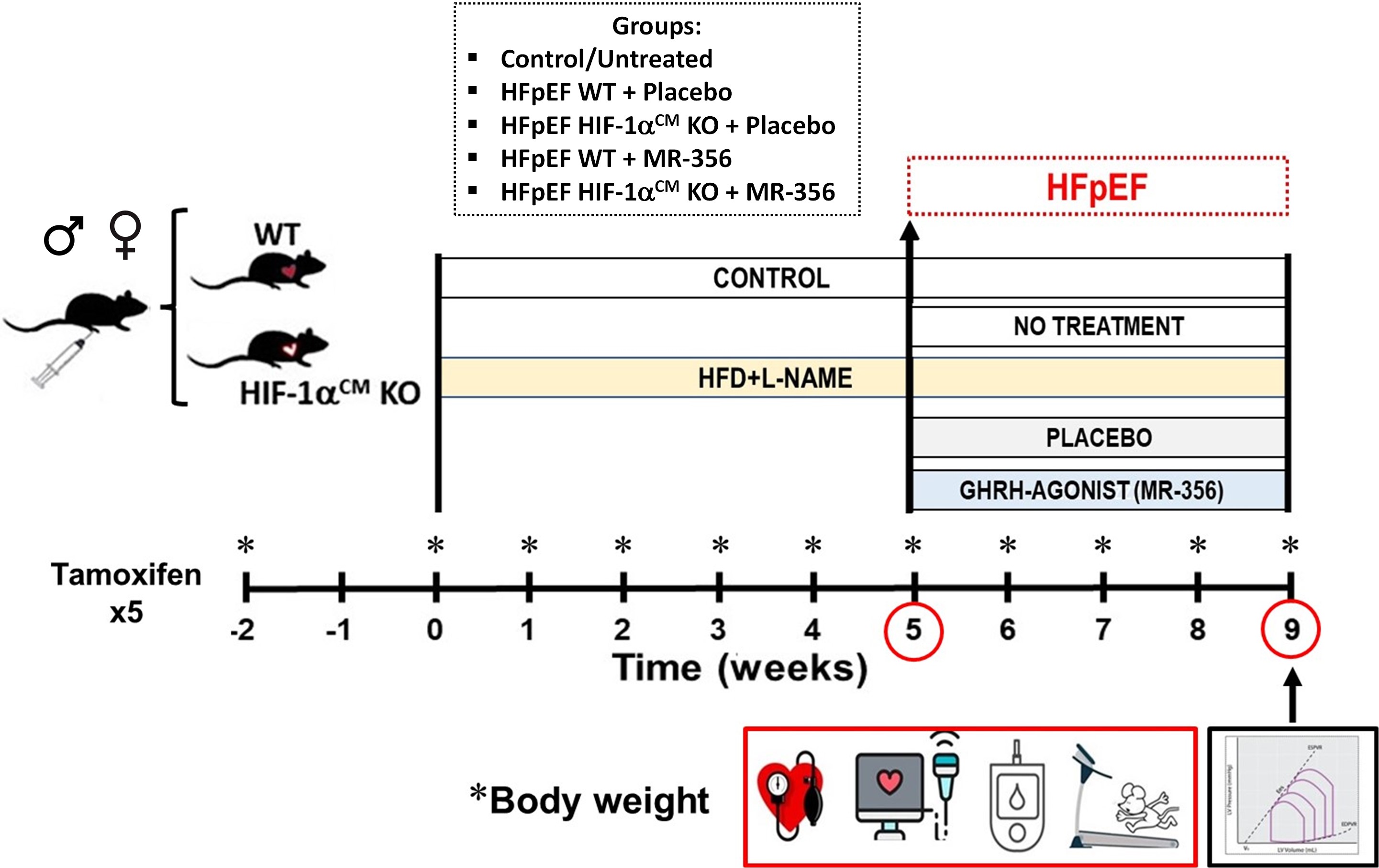

### Supplemental Figure S2

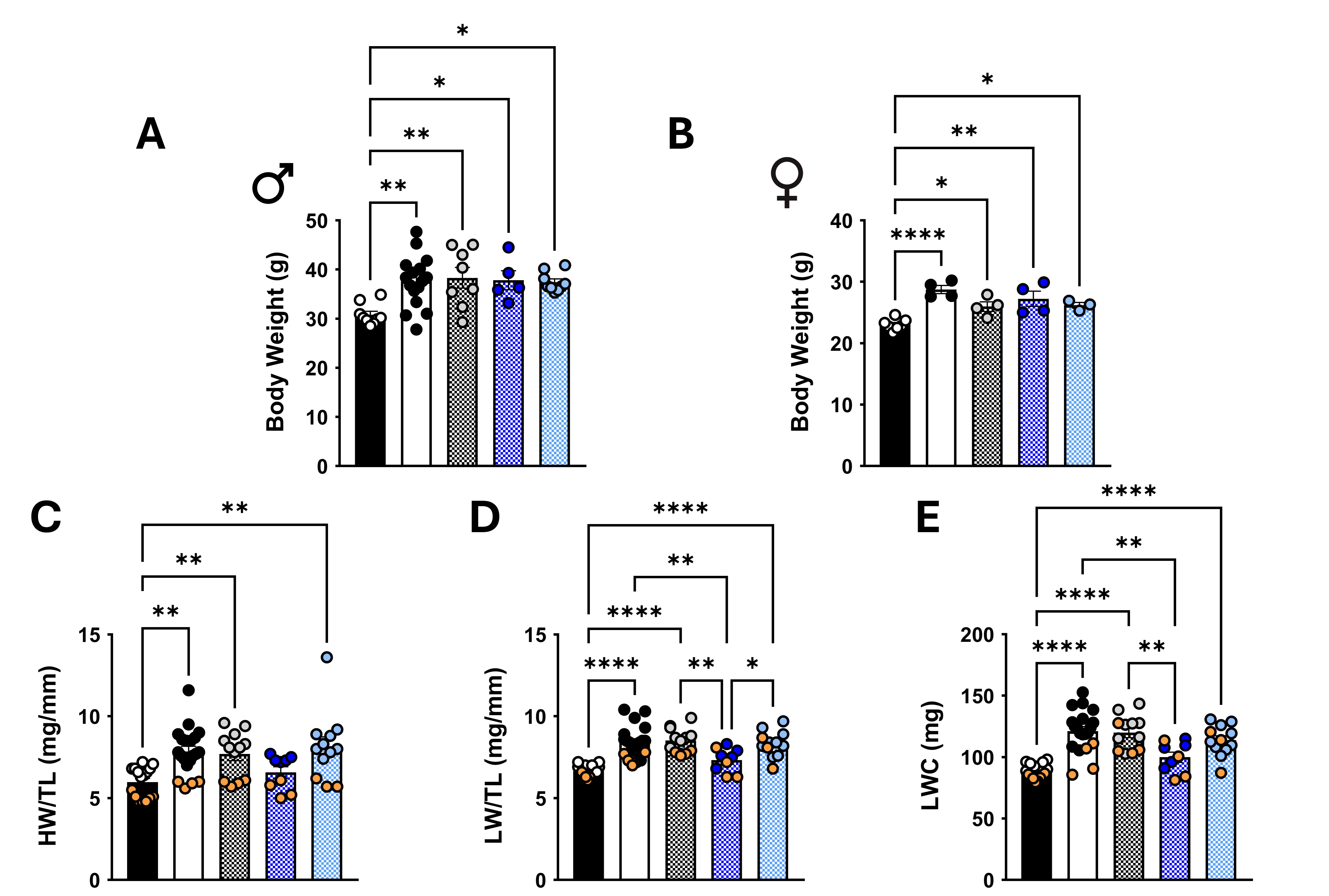
