## Supplemental Table S1 for "Cardiomyocyte-specific expression of HIF-1α mediates the cardioprotective effects of Growth Hormone Releasing Hormone (GHRH)"

### Supplemental Table S1. Hemodynamic parameters

| Parameters | CONTROL | HFpEF | | | | *P value* |
|---|---|---|---|---|---|---|
| | | PLACEBO | | GHRH-AGONIST MR-356 | | |
| | (N=16) | WT (N=18) | HIF-1a$^{CM}$KO (N=10) | WT (N=10) | HIF-1a$^{CM}$KO (N=10) | |
| **Heart rate (bpm)** | 500±14 | 490±9 | 523±11 | 479±16 | 502±16 | ns |
| ***Integrated Performance*** | | | | | | |
| **SW (mmHg x ml)** | 2028 ± 166.9 | 1969 ± 132.2 | 2262 ± 184.6 | 2468 ± 194.2 | 2172± 179.1 | ns |
| **SV (ml)** | 26.7 ± 1.5 | 27.9 ± 4.2 | 24.71 ± 1.5 | 26.7 ± 1.8 | 24.4 ± 2.5 | ns |
| **CO (ml/min)** | 13.9 ± 0.7 | 11.6 ± 1.6 | 12.5 ± 0.7 | 13.95 ± 0.5 | 12.9 ± 0.1 | ns |
| **Ea/Ees** | 1.2 ± 0.5 | 1.3 ± 0.2 | 1.7 ± 0.1 | 1.4 ± 0.2 | 1.3 ± 0.8 | ns |
| ***Afterload*** | | | | | | |
| **ESP (mmHg)** | 85.7 ± 1.2 | 104.0± 2.1 * | 110.9 ± 5.7 * | 105.6 ± 4.8 * | 107.5 ± 5.7 * | * p<0.05 vs. Control |
| **Ea (mmHg/ml)** | 2.8 ± 0.1 | 3.7 ± 0.5 | 3.9 ± 0.6 | 3.6 ± 0.3 | 3.8 ± 0.3 | ns |
| ***Preload*** | | | | | | |
| **EDP (mmHg)** | 5.0 ± 0.3 | 11.8 ± 1.1 § | 14.2 ± 2.0 § | 6.2 ± 0.8 † | 8.7 ± 0.8 * | * p<0.05, § p<0.0001 vs. Control<br>† p< 0.01 vs. WT Placebo and HIF-1a$^{CM}$KO Placebo |
| **EDV (ml)** | 43.3 ± 2.7 | 42.5 ± 1.9 | 48.6± 1.7 | 43.6 ± 2.5 | 49.8 ± 2.1 | ns |
| ***Contractility*** | | | | | | |
| **dP/dT$_{max}$ (mmHg/s)** | 7241 ± 395 | 7108 ± 426 | 7982 ± 677 | 7349 ± 589 | 8086 ± 722 | ns |
| **Ees (mmHg/ml)** | 2.3 ± 0.3 | 3.0 ± 0.4 | 1.8 ± 0.9 | 3.6 ± 0.5 | 3.7 ± 0.9 | ns |
| **PRSW (mmHg)** | 54.4 ± 2.8 | 57.5 ± 1.3 | 50.3 ± 2.2 | 59.9 ± 3.7 | 58.7 ± 6.2 | ns |
| ***Lusitropy*** | | | | | | |
| **dP/dT$_{min}$ (mmHg/s)** | -7163 ± 395 | -7589 ± 690 | -7078 ± 566 | -7786 ± 567 | -8135 ± 522 | ns |
| **EDPVR (exponential)** | 0.01 ± 0.002 | 0.04 ± 0.002 § | 0.04 ± 0.005 § | 0.02 ± 0.002 *† | 0.03 ± 0.003 † | † p< 0.01 and § p<0.0001 vs. Control<br>‡ p<0.001 vs. HIF-1a$^{CM}$KO MR-356<br>† p< 0.01 vs. WT Placebo and HIF-1a$^{CM}$KO Placebo<br>*p<0.005 vs. HIF-1a$^{CM}$KO MR-356 |

Heart rate (HR), stroke work (SW), stroke volume (SV), cardiac output (CO), arterial elastance (Ea) and slope of end-systolic pressure volume relationship (Ees) ratio (Ea/Ees), left ventricle end-systolic pressure (ESP), arterial elastance (Ea), end-diastolic pressure (EDP), left ventricle end-diastolic volume (EDV), maximal rate of pressure rise (dP/dTmax), end-systolic pressure volume relationship [ESPVR](Ees), preload recruitable stroke work (PRSW), maximal rate of pressure decline (dP/dTmin), end-diastolic pressure volume relationship (EDPVR).

All data are presented as mean ± SE. One-way ANOVA followed by Tukey's multiple comparisons test or Kruskal-Wallis followed by Dunn's multiple comparisons test.
