## Supplemental Table S2 for "Cardiomyocyte-specific expression of HIF-1α mediates the cardioprotective effects of Growth Hormone Releasing Hormone (GHRH)"

### Supplemental Table S2. Gravimetric data.

| MALE Parameters | CONTROL | HFpEF | | | | P value |
|---|---|---|---|---|---|---|
| | | PLACEBO | | GHRH-AGONIST (MR-356) | | |
| | (N=9) | WT (N=16) | HIF-1α$^{CM}$KO (N=8) | WT (N=5) | HIF-1α$^{CM}$KO (N=9) | |
| BW (g) | 30.8±0.7 | 37.6±1.3 † | 38.3±2.1 † | 37.8±1.9 * | 37.4±0.6 * | * p< 0.05, † p<0.01 vs. Control |
| TL (mm) | 18.6±0.1 | 18.5±0.1 | 18.7±0.1 | 18.7±0.2 | 18.3±0.1 | ns |
| HW (mg) | 125.4±2.0 | 154.3±5.2 ‡ | 161.1±4.9 ‡ | 138.3±1.0 | 162.3±11.2 ‡ | ‡ p<0.001 vs. Control |
| LW (mg) | 128.3±1.8 | 162.0±4.1 § | 163.8±5.1 § | 142.4±4.5 * | 154.9±4.2 † | † p<0.01, § p<0.0001 vs. Control<br>* p<0.05 vs. WT Placebo |
| HW/TL (mg/mm) | 6.7±0.1 | 8.3±0.3 † | 8.6±0.2 † | 7.4±0.1 | 8.9±0.6 † | † p<0.001 vs. Control |
| LW/TL (mg/mm) | 6.9±0.1 | 8.6±0.2 § | 8.7±0.2 § | 7.6±0.2 | 8.5±0.2 ‡ | ‡ p<0.001, § p<0.0001 vs. Control |
| LWC (mg) | 92.6±1.4 | 126.8±3.2 § | 124.7±4.6 § | 103.9±4.4 *† | 115.9±3.6 ‡ | ‡ p<0.001, § p<0.0001 vs. Control<br>† p< 0.01 vs. WT Placebo<br>*p<0.05 vs. HIF-1α$^{CM}$KO MR-356 |

| FEMALE Parameters | CONTROL | HFpEF | | | | P value |
|---|---|---|---|---|---|---|
| | | PLACEBO | | GHRH-AGONIST (MR-356) | | |
| | (N=7) | WT (N=4) | HIF-1α$^{CM}$KO (N=4) | WT (N=4) | HIF-1α$^{CM}$KO (N=3) | |
| BW (g) | 23.0±0.3 | 28.7±0.6 § | 26.0±0.8 * | 27.2±1.2 † | 26.2±0.5 * | * p< 0.05, † p<0.01 , § p<0.0001vs. Control |
| TL (mm) | 18.1±0.1 | 18.0±0.1 | 18.0±0.06 | 18.1±0.06 | 18.0±0.2 | ns |
| HW (mg) | 91.1±2.1 | 105.8±1.5 | 106.3±1.5 * | 100.1±4.1 | 106.3±1.5 | * p< 0.05 vs. Control |
| LW (mg) | 115.5±1.2 | 131.6±6.7 | 145.0±6.6 | 125.9±7.5 | 140.4±10.2 | ns |
| HW/TL (mg/mm) | 5.0±0.1 | 5.9±0.1 | 5.9±0.1 * | 5.5±0.2 | 5.9±0.2 | * p< 0.05 vs. Control |
| LW/TL (mg/mm) | 6.4±0.07 | 7.6±0.03 | 8.1±0.4 * | 7.0±0.4 | 7.9±0.26 | * p< 0.05 vs. Control |
| LWC (mg) | 83.1±1.0 | 98.6±6.2 | 110.5±5.8 * | 94.9±7.5 | 107.3±10.2 | * p< 0.05 vs. Control |

Morphometric data are normalized to tibia length (TL). Values are expressed as means ± standard error of mean (SEM). Male (M) and female (F). Body weight (BW), tibia length (TL), heart weight (HW), lung weight (LW), ratio of heart weight to tibia length (HW/TL), ratio of lung weight to tibia length (LW/TL), lung water content (LWC).

One-way ANOVA followed by Tukey’s multiple comparisons test or Kruskal-Wallis followed by Dunn’s multiple comparisons test.
